# A catalog of transcription start sites across 115 human tissue and cell types

**DOI:** 10.1101/2021.05.12.443890

**Authors:** Jill E. Moore, Xiao-Ou Zhang, Shaimae I. Elhajjajy, Kaili Fan, Fairlie Reese, Ali Mortazavi, Zhiping Weng

## Abstract

Accurate transcription start site (TSS) annotations are essential for understanding transcriptional regulation and its role in human disease. Gene collections such as GENCODE contain annotations for tens of thousands of TSSs, but not all of these annotations are experimentally validated nor do they contain information on cell type-specific usage. Therefore, we sought to generate a collection of experimentally validated TSSs by integrating *RNA Annotation and Mapping of Promoters for the Analysis of Gene Expression* (RAMPAGE) data from 115 cell and tissue types, which resulted in a collection of approximately 50 thousand representative RAMPAGE peaks. These peaks were primarily proximal to GENCODE-annotated TSSs and were concordant with other transcription assays. Because RAMPAGE uses paired-end reads, we were then able to connect peaks to transcripts by analyzing the genomic positions of the 3’ ends of read mates. Using this paired-end information, we classified the vast majority (37 thousand) of our RAMPAGE peaks as verified TSSs, updating TSS annotations for 20% of GENCODE genes. We also found that these updated TSS annotations were supported by epigenomic and other transcriptomic datasets. To demonstrate the utility of this RAMPAGE rPeak collection, we intersected it with the NHGRI/EBI GWAS catalog and identified new candidate GWAS genes. Overall, our work demonstrates the importance of integrating experimental data to further refine TSS annotations and provides a valuable resource for the biological community.

## Introduction

Accurate maps of genes and their transcription start sites (TSSs) are essential for studying gene regulation and determining the impact of genetic variation. While gene and transcript annotations have improved substantially over the years, benefiting from advances in experimental and computational technologies, accurate, cell type-specific annotations are far from complete. Efforts such as the GENCODE project (Frankish et al. 2019) have generated detailed annotations for over 60 thousand genes and 100 thousand transcripts across the human genome. These widely used annotations combine transcriptomic, proteomic, and homology evidence through manual curation and automated computational pipelines. However, these annotations are built in a cell type-agnostic manner; they represent the collective transcriptomic landscape across thousands of unique cell and tissue types. Therefore, it is difficult to know which transcripts are actively transcribed in a particular cell or tissue type and, by extension, which regulatory elements and genetic variants may impact gene expression.

While public RNA-seq data are accumulating across a wide array of tissues and cells types, many of which are from coordinated efforts such as the Genotype-Tissue Expression (GTEx) (GTEx Consortium 2020) and Encyclopedia of DNA Elements (ENCODE) (ENCODE Project Consortium et al. 2020) projects, these experiments are not optimal for annotating specific transcripts and their start sites. Most RNA-seq protocols perform short-read sequencing, which can accurately quantify gene expression levels but are unable to fully delineate transcript isoforms nor precisely map the 5’ ends of transcripts. Therefore, assays that target and preserve 5’ ends, such as the *cap analysis gene expression* (CAGE) assay (Kodzius et al. 2006), are preferred for TSS identification. The FANTOM consortium generated a TSS catalog across the human genome by integrating thousands of CAGE experiments (FANTOM Consortium and the RIKEN PMI and CLST (DGT) et al. 2014). However, CAGE uses short, single-end reads, which have low mappability and can not connect TSSs to their downstream transcripts. To overcome these limitations, Gingeras and colleagues developed the *RNA Annotation and Mapping of Promoters for the Analysis of Gene Expression* (RAMPAGE) assay (Batut et al. 2013), which captures the 5’ end of capped RNAs using paired-end reads to enable more accurate genomic mapping and transcript characterization. The Gingeras lab generated both RAMPAGE and RNA-seq data for more than one-hundred human samples during the ENCODE Project (ENCODE Project Consortium et al. 2020).

Here, we integrated 115 high-quality ENCODE RAMPAGE experiments to identify 52,546 representative RAMPAGE peaks (rPeaks), a curated collection of TSSs and their activities across the 115 human samples. These rPeaks are supported by other transcription assays including CAGE, long-read RNA-seq using the PacBio platform, and high-resolution nuclear run-on of capped transcripts (GRO-cap) (Core et al. 2014). Using paired-end RAMPAGE reads, we linked the majority of rPeaks to annotated genes and identified TSSs of unannotated spliced transcripts. These verified rPeaks were more enriched for transcriptomic and epigenomic features than GENCODE TSSs for the same genes not supported by RAMPAGE. Finally, we used this collection of rPeaks to annotate human variants associated with genome-wide association studies (GWAS) and identify novel phenotype-associated genes. Overall, our TSS collection complements existing gene annotations and demonstrates the utility of cell type-specific TSS annotations in integrative analyses.

## Results

### Curation of 52,546 representative RAMPAGE peaks

We curated 115 high-quality RAMPAGE experiments (**Supplemental Table S1a**) from ENCODE to generate our collection of representative RAMPAGE peaks (rPeaks) (**Fig. 1a**). These RAMPAGE experiments spanned 87 tissues and 28 cell types from a variety of biological contexts. We called peaks in individual RAMPAGE experiments as previously described (Zhang et al. 2019), identifying three components for each peak: (1) a full peak; (2) a high density region in the peak that accounts for 70% of the peak’s total RAMPAGE signal; and (3) a summit, which is the genomic position with the highest signal. Given that the RAMPAGE assay enriched for reads at the 5’ ends of transcripts, we filtered out the small subset of RAMPAGE peaks that had higher RNA-seq signals than RAMPAGE signals in the matched biosample (see *Methods,* **Supplemental Fig. S1a**), retaining about ten thousand peaks per experiment (**Supplemental Table S1a**). We then clustered overlapping peaks across the 115 experiments for two genomic strands separately and selected a representative peak (rPeak) for each cluster with the highest Reads Per Kilobase, per Million mapped reads (RPKM, **Fig. 1a**). Additional filtering was performed to remove low signal, single-experiment rPeaks which were likely false positives; in total, we arrived at 52,546 rPeaks (**Supplemental Table S1b**). The full rPeaks and their high-density regions occupy 0.23% and 0.09% of the human genome, having median widths of 121 and 43 nucleotides (nts), respectively (**Supplemental Fig. S1b, c**).

**Figure 1.**
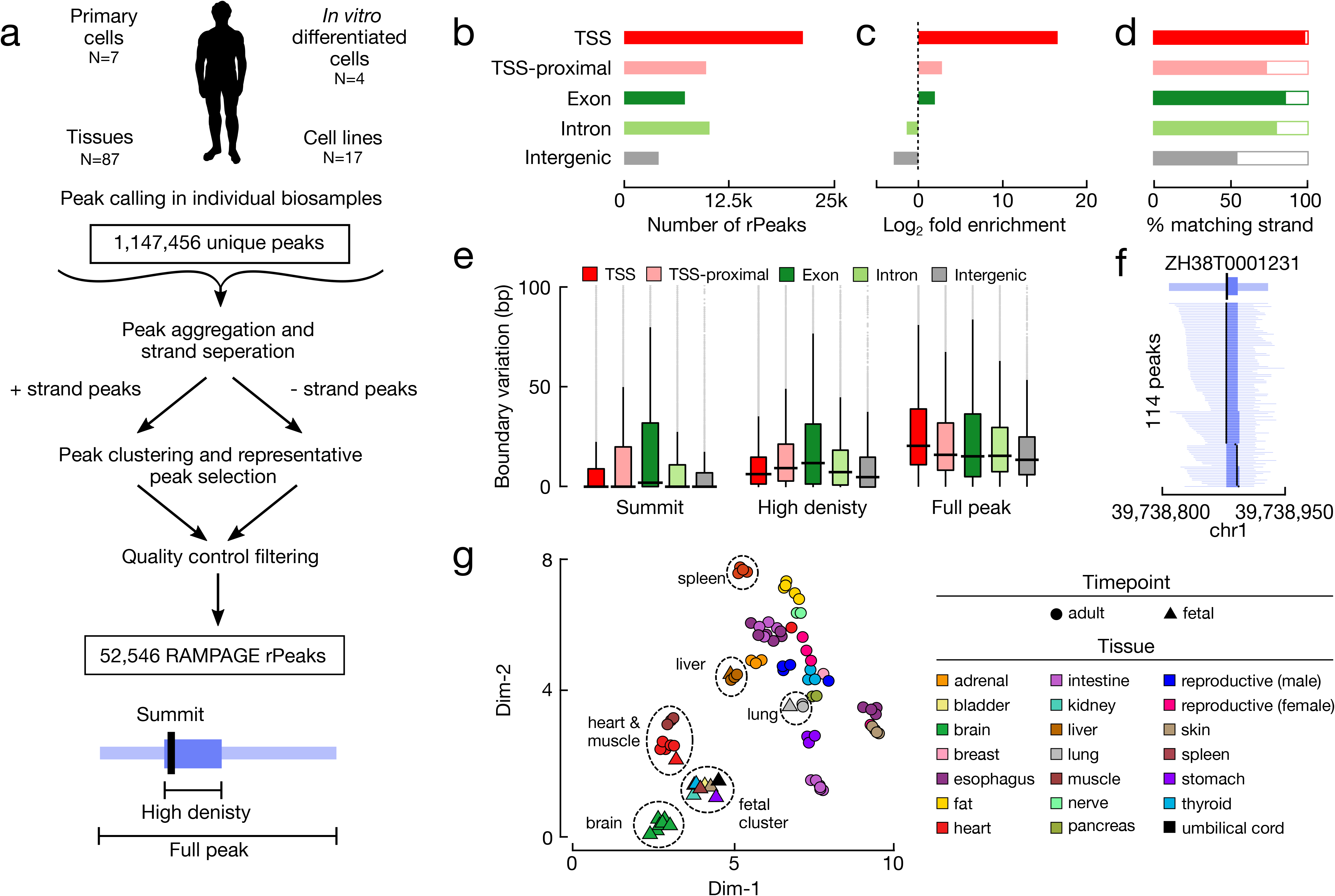
Curating a collection of representative RAMPAGE peaks (rPeaks) across 115 biosamples. **a,** Workflow for curating RAMPAGE rPeaks. First, we called peaks in individual RAMPAGE datasets across 115 cell types and tissues. We then pooled these peaks (N=1,147,456) and separated them by genomic strand. We clustered overlapping peaks on the same strand, selected the peak with the highest RAMPAGE signal (i.e., the rPeak) to represent each cluster, removed all the peaks overlapping the rPeak from the pool, and performed clustering on the remaining peaks. We repeated this process iteratively until all peaks were accounted for by rPeaks. We performed additional filtering using RNA-seq data, removing peaks that had a higher RNA-seq signal than RAMPAGE signal, finally arriving at 52,546 rPeaks. **b,** Bar plots showing the number of RAMPAGE rPeaks stratified into distinct sets by genome context: overlapping GENCODE V31 TSSs (red), proximal (± 500 bp) to TSSs (pink), overlapping exons (dark green), overlapping introns (light green), and intergenic (gray). **c**, Bar plots showing the fold enrichment for the number of genomic positions covered by rPeaks over the footprints of the genomic contexts in **b**. **d**, Bar plots showing the percentage of rPeaks in each genomic context as in **b** that are on the same strand as their overlapping TSS, gene (exon and intron), or nearest gene (TSS-proximal and intergenic). **e**, Box plots displaying the variation in the positions of rPeak summits (left), high-density region boundaries (middle), and full peak boundaries (right), stratified by the genomic contexts as in **b**. **f**, An example TSS-overlapping rPeak ZH38T000123 from K562 cells and the RAMPAGE peaks it represents in 113 other biosamples. For each peak, the full width is denoted in light blue, high-density regions in blue, and summit in black. **g**, Scatterplot displaying a two-dimensional Uniform Manifold Approximation and Projection (UMAP) embedding of 87 tissue samples using RAMPAGE signal across all rPeaks as input features. Circles denote adult tissues and triangles denote fetal tissues. Markers are colored by tissue of origin as defined in the legend.

The majority (59%) of rPeaks either overlapped or were proximal to (± 500 bp but did not overlap) a GENCODE-annotated TSS (GENCODE 31 basic TSSs; **Fig. 1b**). The remaining rPeaks overlapped exonic, intronic, or intergenic regions (14%, 18%, and 9% of rPeaks, respectively). We used these genomic contexts (TSS, TSS-proximal, exon, intron, and intergenic) throughout our analyses. As expected, RAMPAGE rPeaks were highly enriched for annotated TSSs and depleted in intergenic regions compared to the genomic footprints of these contexts (Chi-square test, *p* < 1 × 10^-300^, **Fig. 1c**). Additionally, TSS, TSS-proximal, exonic, and intronic rPeaks had higher strand concordance than intergenic rPeaks, meaning they were more likely to fall on the same stand as their overlapping or closest gene (**Fig. 1d**). This finding suggests that intergenic rPeaks likely resulted from misannotated or novel TSSs or from transcription at regulatory elements such as enhancer RNAs (eRNAs).

Next, we analyzed the ability of each rPeak to accurately represent their underlying clusters. When we analyzed the range of biosample activities of the RAMPAGE peak clusters, we observed a bimodal distribution (**Supplemental Fig. S1d**, **Supplemental Table S1b**), indicating that some rPeaks represent peaks from many RAMPAGE experiments, while others represent only a few. The rPeaks in different genomic contexts differ greatly in this regard. TSS rPeaks represent peaks from 26 experiments on average, much higher than other rPeaks (pairwise Fisher’s exact test, *p* < 1 × 10^-300^, **Supplemental Fig. S1e**). For the vast majority of rPeaks, their summits were at nearly identical positions to the peaks they represented (we excluded the peak that was chosen as the rPeak for this analysis), with a difference in median of 0 bp across experiments (**Fig. 1e**). We observed more variability in the boundaries of the high-density regions and full peaks, with a difference in median of 8 and 18 bp, respectively, which is still relatively small on a genome-wide scale. As an illustrative example, the rPeak (ZH38T0001231) that overlapped a TSS of the *PPIE* gene represented RAMPAGE peaks from 114 experiments precisely at their summits and high-density regions (**Fig. 1f, Supplemental Table S1c**).

To assess the biological spectrum of rPeak activity, we performed dimensionality reduction with UMAP for all tissue samples (N = 87), using RAMPAGE signal profiles across the rPeaks in these samples (**Fig. 1g**). Similar tissues generally clustered together, e.g., brain, heart and leg muscle, and gastrointestinal tissues, respectively. Some fetal tissues clustered with their corresponding adult tissues, including heart, liver, and lung. However, fetal thyroid and stomach tissues clustered exclusively with other fetal tissue samples, suggesting that these fetal samples share developmental transcriptional patterns at the surveyed life stages. We observed similar patterns using all 115 biosamples, albeit with tissues clustering separately from primary cells and cell lines (**Supplemental Fig. S1f**). Overall, these results indicate that our RAMPAGE rPeaks are a unified set of transcriptional sites that enables systematic investigations into the transcriptional landscape across multiple biosamples.

### RAMPAGE rPeaks are concordant with other transcription start site annotations

To evaluate the accuracy and comprehensiveness of our RAMPAGE rPeaks collection, we compared it with other collections of transcription start site annotations. The largest and most biologically diverse of these collections is the atlas of CAGE peaks generated by the FANTOM5 consortium, which comprises 209,911 peaks annotated across 1,816 experiments (FANTOM Consortium and the RIKEN PMI and CLST (DGT) et al. 2014; Abugessaisa et al. 2017). Approximately two-thirds of our rPeaks overlapped a CAGE peak, while only one-third of the CAGE peaks overlapped an rPeak (**Fig. 2a**). Stratified by genomic context, the CAGE-overlapping rPeaks were more likely to be at GENCODE TSSs or TSS-proximal loci and less likely to be at exonic, intergenic, or intronic loci (Chi-square test, *p* < 1.0 x 10^-300^, **Fig. 2b**).

**Figure 2.**
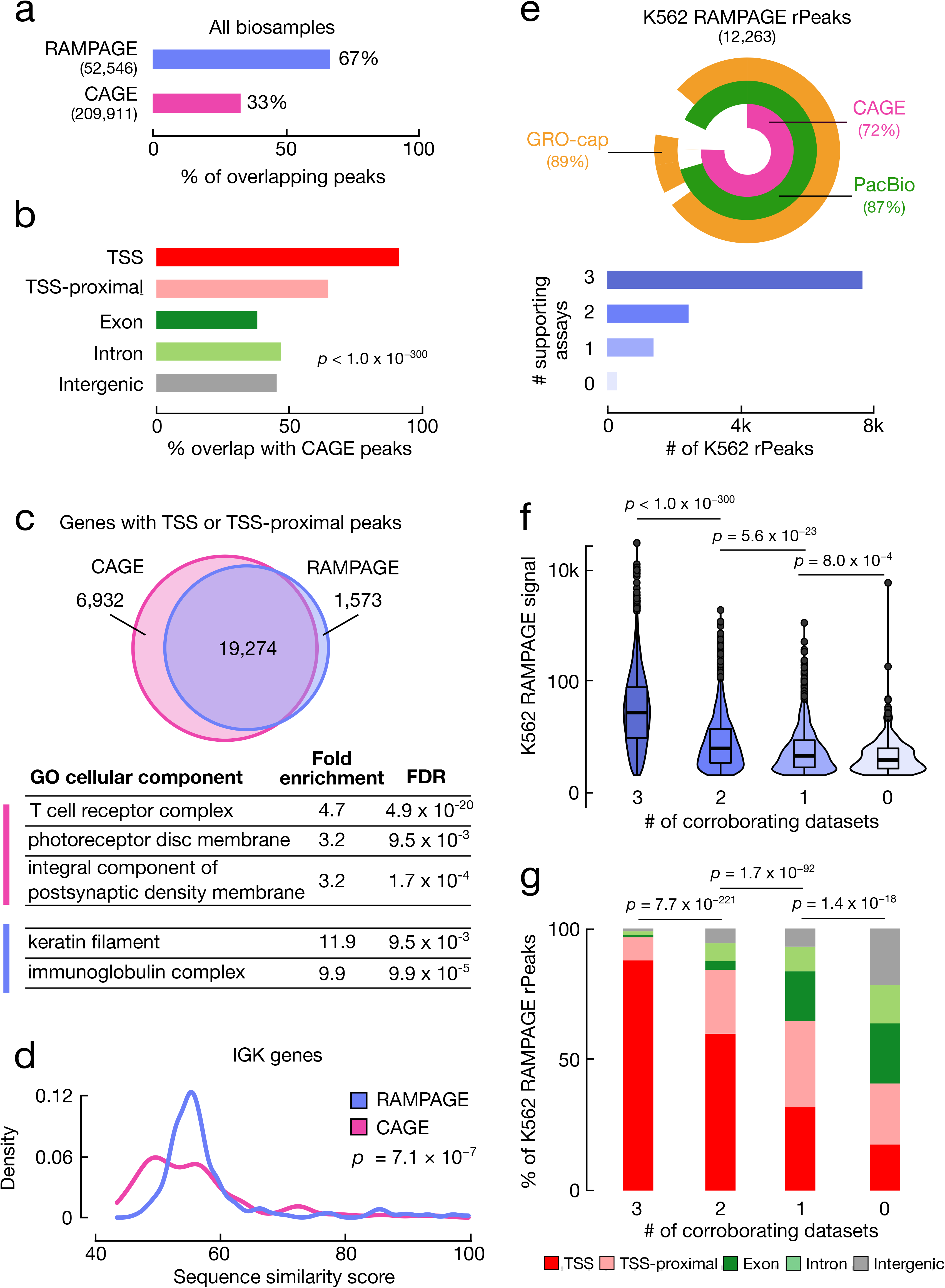
RAMPAGE rPeaks are concordant with other transcriptome annotations. **a,** Bar graph showing the percentage of RAMPAGE rPeaks that overlap CAGE peaks (purple) and the percentage of CAGE peaks that overlap RAMPAGE rPeaks (pink). **b,** Bar graph showing the percentages of CAGE-overlapping RAMPAGE peaks in specific genomic contexts as defined in Fig. 1b. c, Venn diagram depicting the overlap of genes whose TSSs have at least one peak within 500 bp between the sets of CAGE peaks (pink) and RAMPAGE rPeaks (purple). Below are representative gene ontology terms (cellular component) enriched in CAGE-only genes (pink) or RAMPAGE-only genes (purple). A full list of enriched terms can be found in **Supplemental Table S2**. **d,** Density plot showing the distributions of the similarity scores for sequences surrounding the TSSs of immunoglobulin kappa (IGK) genes supported by only RAMPAGE peaks (purple) or only CAGE peaks (pink). Sequence similarity was calculated as the maximal score of all pairwise local alignments. *P-*value corresponds to a two-sided Wilcoxon test. **e,** (top) VennPie diagram with concentric circles displaying K562 RAMPAGE rPeaks that overlap K562 CAGE peaks (pink) or PacBio 5’ ends (green), or have high GRO-seq signals (orange). The overall percentages are shown in parentheses. (bottom) Bar plot with the number of K562 rPeaks stratified by the number of supporting transcriptomic assays in the above VennPie. **f,** Violin-boxplot showing the distributions of the K562 RAMPAGE signal of rPeaks stratified by the number of supporting assays as defined in **e.** *P*-values correspond to two-sided pairwise Wilcoxon tests with FDR correction. **g,** Stacked bar graphs showing the percentage of K562 rPeaks belonging to each genomic context (TSS: red, TSS-proximal: pink, exon: dark green, intron: light green, intergenic: gray) stratified by the number of supporting assays as defined in **e.** *P*-values correspond to Chi-square tests.

Though a majority of TSS and TSS-proximal rPeaks were shared between RAMPAGE and CAGE, we identified sets of genes with TSSs that were exclusively proximal to CAGE peaks (CAGE-only genes, **Supplemental Table S2a**) or were exclusively proximal to RAMPAGE rPeaks (RAMPAGE-only genes, **Supplemental Table S2b**). Gene ontology analysis revealed that the 6,932 CAGE-only genes were enriched in terms such as *T cell receptor complex* and *photoreceptor disc membrane* (**Fig. 2c, Supplemental Table S2c**). Enrichment in these terms is not surprising as the corresponding biosamples-T cells and eye tissues-were assayed by CAGE but not RAMPAGE. Additional enrichment for neuronal terms such as *integral components of postsynaptic density membranes* was unexpected as we integrated RAMPAGE data from seven fetal brain samples and *in vitro* differentiated neurons. Alternatively, the 1,573 RAMPAGE-only genes were primarily enriched for two terms, immunoglobulin production and keratinization (**Fig. 2c, Supplemental Table S2d**), due to an abundance of immunoglobulin kappa (*IGK*) genes and keratin-associated protein (*KRTAP*) genes, respectively. While we observed genes from these families in both the RAMPAGE-only and CAGE-only gene sets, the sequences flanking the TSSs of RAMPAGE-only *IGK* and *KRTAP* genes shared higher sequence similarity than the corresponding CAGE-only genes (Wilcoxon tests, *p* = 7.1 x 10^-7^ and 4.7 x 10^-5^, respectively, **Fig. 2d, Supplemental Fig. S2a**). We hypothesize that the 101 nt, paired-end RAMPAGE reads can uniquely map to genomic regions sharing higher sequence similarity better than the 36 nt, single-end CAGE reads, allowing us to identify rPeaks in more paralogs of these two gene families. In conclusion, the primary differences in the gene coverage by RAMPAGE rPeaks and CAGE peaks largely reflect differences in their biosample collections and assay read length.

To avoid coverage differences due to biosample composition, we directly compared RAMPAGE rPeaks and CAGE peaks active in K562 and GM12878 cell lines-biosamples used in both peak collections-along with GRO-cap (Core et al. 2014) and PacBio long-read RNA-seq data (Wyman et al.) in the respective cell lines. Generally, the four sets of TSS annotations were highly concordant, with the majority of RAMPAGE rPeaks, CAGE peaks, and PacBio 5’ ends overlapping one another and containing high GRO-cap signals (**Supplemental Fig. S2b**). In total, 98% of K562 and GM12878 rPeaks were substantiated by overlaps with at least one other transcriptome annotation, and a majority of the rPeaks-65% in K562 and 74% in GM12878-was supported by all of the other assays (**Fig. 2e** and **Supplemental Fig. S2c)**. These supported rPeaks had higher RAMPAGE signals (Pairwise Wilcoxon test, *p* < 1.0 × 10^-300^, **Fig. 2f, Supplemental Fig. S2d**) and were more likely to overlap TSSs (Chi-square test, *p* = 2.5 × 10^-259^, **Fig. 2g, Supplemental Fig. S2e**) than rPeaks supported by fewer or no other assays. We also analyzed the number of genes with TSSs supported by RAMPAGE, CAGE or PacBio data, and observed that RAMPAGE rPeaks identified an average of 17% fewer genes than CAGE and PacBio (**Supplemental Fig. S2f, g**). On the other hand, 96% of RAMPAGE genes were supported by another assay compared to 90% of CAGE genes and 88% of PacBio genes. Thus, our rPeak approach slightly compromises recall for better precision.

Stratified by genomic context, TSS rPeaks had the highest levels of GRO-cap signal and overlapped the greatest number of PacBio 5’-ends, followed by TSS-proximal, intronic and intergenic rPeaks (**Supplemental Fig. S2h-j**). In contrast, exonic rPeaks consistently had the lowest levels of GRO-cap signal, and though they overlapped a moderate number of PacBio 5’-ends (**Supplemental Fig. S2i**), these PacBio reads were significantly shorter than those overlapping other rPeak classes (pairwise Wilcoxon test with FDR correction, *p* < 1.0 × 10^-16^, **Supplemental Fig. S2j**). These results suggest that many of the exonic rPeaks likely are not TSSs and may arise from mRNA recapping (Trotman and Schoenberg 2019).

Only 2% of K562 and GM12878 RAMPAGE rPeaks were not supported by one of the other assays (N = 301 and 212, respectively). These peaks had the lowest RAMPAGE signals and were more likely to overlap exons, introns, and intergenic regions (**Fig. 2f-g, Supplemental Fig. S2d-e)**. These weaker transcription sites are not as reproducible across assays or may be false positives. Overall, our results demonstrate that our set of RAMPAGE rPeaks are highly concordant with other transcriptome annotations and likely represent a conservative set of transcription start sites.

### Three-quarters of RAMPAGE rPeaks are assigned to genes via spliced transcripts

One advantage of the RAMPAGE assay is that it produces paired-end reads, which not only result in more accurately mapped fragments but also have the ability to assign rPeaks to genes. rPeaks are derived from the 5’ ends of RAMPAGE read pairs, and by analyzing the genomic positions of 3’ ends of the read pairs, we can attempt to link rPeaks to downstream transcripts and consequently assign rPeaks to genes. Such a process could go down one of two general paths (**Fig. 3a**). If the generated transcript is spliced (e.g., mRNAs and lncRNAs), the 3’ end of the read pair would map to an exon that is thousands of base pairs downstream of the rPeak, and we can assign the rPeak to the gene that the exon belongs to. However, if the generated transcript is unspliced (e.g. pre-mRNAs, single-exon transcripts, small RNAs, and most eRNAs), the 3’ end will map less than 1 kilobases (kb) downstream (the maximum selected fragment size for the RAMPAGE assay; median = 335 nt) and we cannot confidently assign the rPeak to a specific gene.

**Figure 3.**
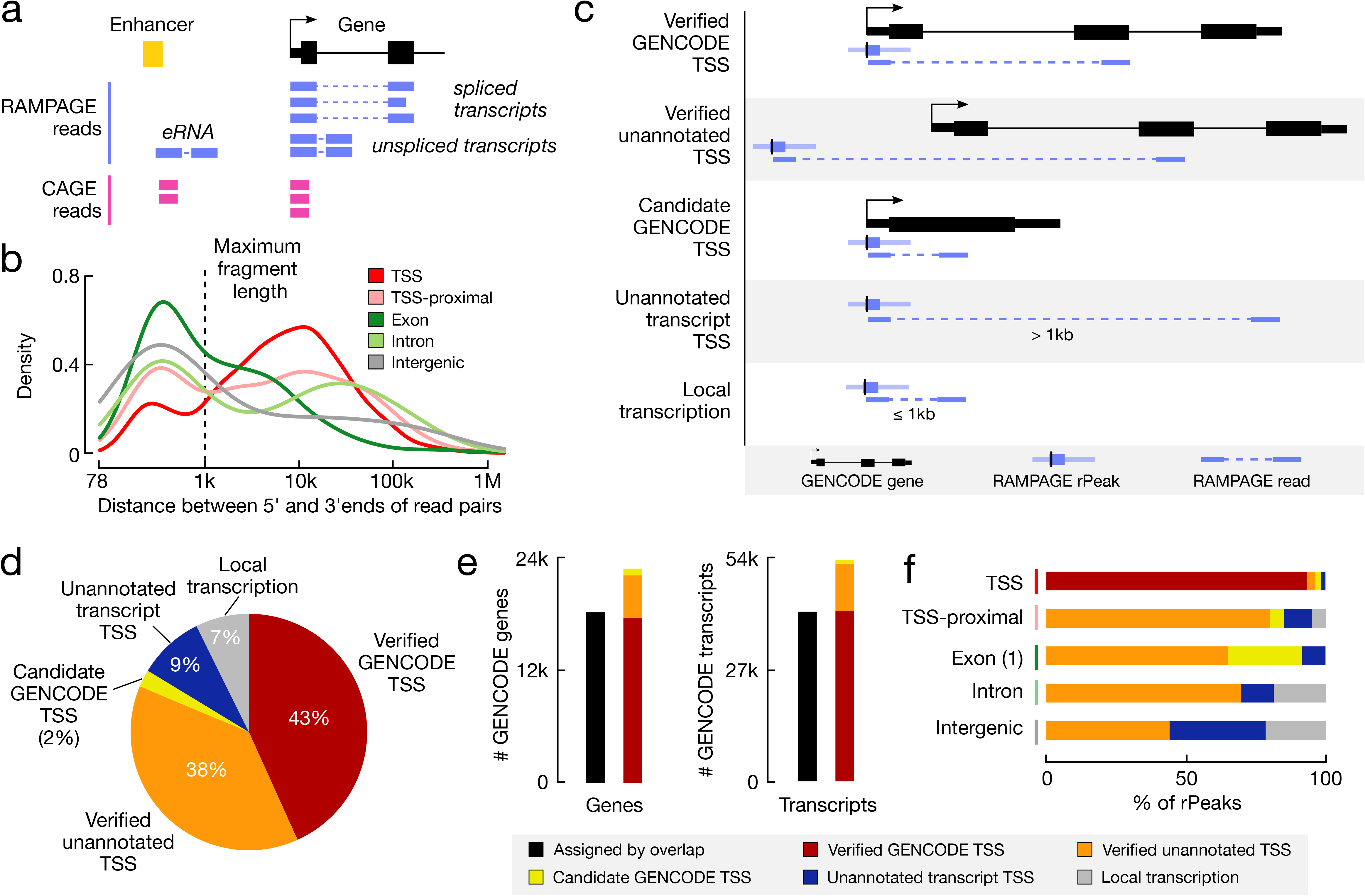
Assigning RAMPAGE rPeaks to genes using paired-end reads. **a,** Schematic demonstrating how paired-end RAMPAGE reads (purple) can distinguish between spliced and unspliced transcripts, unlike single-end CAGE reads (pink). **b,** Density plot of the distances between the 5’ and 3’ ends of RAMPAGE read pairs, stratified by rPeak genomic context. The maximum fragment length (1 kb) is shown by the dashed line. **c,** Schematic depicting the paired-end read method for linking RAMPAGE rPeaks with genes and the resulting five categories. **d,** Pie chart displaying the percentage of RAMPAGE rPeaks classified as the five categories in **c**: verified GENCODE TSSs (red), verified unannotated TSSs (orange), candidate GENCODE TSSs (yellow), unannotated transcript TSSs (blue), or local transcription (gray). **e.** Bar graphs showing the number of GENCODE genes (left) and transcripts (right) that are accounted for by overlapping RAMPAGE rPeaks (black) vs. the paired-end read method illustrated in **a** and **c** (colors). Bars for the paired-end method are stratified by TSS class (as defined in **c**, **d**). Genes with multiple TSSs were counted only once with the following priority: verified GENCODE TSSs, verified unannotated TSSs, then candidate GENCODE TSSs. **f,** Bar graphs showing the percentage of rPeaks that are classified as verified GENCODE TSSs (red), verified unannotated TSSs (orange), candidate GENCODE TSSs (yellow), unannotated transcript TSSs (blue), or local transcription (gray), stratified by genomic context.

To determine the portions of RAMPAGE reads derived from spliced and unspliced transcripts, we calculated the distance between the 5’ and 3’ ends of the read pairs that support an rPeak (i.e., the 5’ end of the read pair overlaps an rPeak). For TSS-overlapping rPeaks, we observed a bimodal distribution, with 83% of reads from spliced transcripts (distance > 1 kb) and 17% from unspliced transcripts (distance ≤ 1 kb, **Fig. 3b**). For other rPeak classes, we also observed substantial percentages of reads deriving from spliced transcripts (43-68%), suggesting that these rPeaks may correspond to TSSs for misannotated transcripts, novel isoforms, or novel genes. These results indicated that we can use RAMPAGE read pairs from spliced transcripts to assign rPeaks to genes.

We developed a computational pipeline to systematically assign rPeaks to genes (**Supplemental Fig. S3a**). Because of the aforementioned low GRO-cap signal at exonic rPeaks, we excluded all exonic rPeaks that did not overlap the first exon of an annotated transcript (N = 6,709) from this analysis as they likely capture recapping or degradation events rather than sites of transcription. We further discarded four rPeaks whose supporting reads mapped more than 500 kb away. We classified the remaining 45,833 rPeaks into five general categories depending on whether an rPeak overlaps a GENCODE-annotated TSS and whether its supporting reads overlapped a GENCODE-annotated exon, with the most prominent scenarios summarized as follows (**Fig. 3c**) and with details provided in **Supplemental Table S3a**. First, if an rPeak overlaps a GENCODE-annotated TSS and its supporting reads overlap a downstream exon of the same gene, the rPeak is classified as verified GENCODE TSS. Second, if an rPeak does not overlap a GENCODE-annotated TSS but its supporting reads overlap a GENCODE-annotated exon, the rPeak is classified as verified unannotated TSS. Third, if an rPeak overlaps a GENCODE-annotated TSS and its supporting reads overlap the first exon of the gene, and if the gene has only one exon or its first exon is longer than 500 nts, then the rPeak is classified as candidate GENCODE TSS. Fourth, if an rPeak’s supporting reads map to more than 1 kb downstream from the rPeak and do not overlap a GENCODE-annotated exon, the rPeak is deemed to be the TSS of an unannotated transcript. Fifth, if an rPeak’s supporting reads are within 1 kb of the rPeak and the rPeak is not a candidate GENCODE TSS (third category above), then it is deemed to originate from local (i.e., unspliced) transcription. Using our pipeline, we assigned 84% of rPeaks to genes (the first three categories, **Fig. 3d**), which is 4,641 more genes and 12,466 more transcripts than simply using overlaps (**Fig. 3e**). In total, we curated 19,821 verified GENCODE TSSs, 17,447 verified unannotated TSSs, and 1,088 candidate GENCODE TSSs for 22,801 genes and 4,129 TSSs for unannotated transcripts (**Supplemental Table S3a**).

The vast majority of TSS-overlapping rPeaks (19,821 of 21,278, 93%) are verified GENCODE TSSs, indicating that our approach is of high accuracy (**Fig. 3f, Supplemental Table S3b**). These verified GENCODE TSSs amount to 43% of all rPeaks. The next largest category of rPeaks is verified unannotated TSS (38%, N = 17,447, **Supplemental Table S3c**), and these rPeaks are potentially novel TSSs of spliced GENCODE-annotated genes revealed by our collection of RAMPAGE data. Three examples from K562 cells are highlighted (**Supplemental Fig. S3b-d**): exonic rPeak ZH38T0014149, a verified TSS of *ING1,* (**Supplemental Fig. S3b**); intronic rPeak ZH38T0050003, a verified TSS of *GALNT12* (**Supplemental Fig. S3c**); and intergenic rPeak ZH38T0049993, a verified TSS of *NANS* (**Supplemental Fig. S3d)**. These three verified TSSs and their linked transcripts were also supported by PacBio reads in K562 cells.

The candidate GENCODE TSS category of rPeaks (N = 452) constitutes 88% of the TSS-overlapping rPeaks that were not supported by reads from spliced transcripts (N = 511). The GENCODE genes that overlap these rPeaks either have only one exon or have a long (> 500 nts) first exon (**Supplemental Table S3d**). We observed a similar pattern for TSS-proximal and exonic rPeaks not supported by reads from spliced transcripts, though at lower percentages. These rPeaks are most likely the TSSs of the overlapping genes, although our paired-end mapping approach is not able to make the assignment definitively; thus, we assigned them the candidate designation.

The unannotated transcript category includes 4,129 rPeaks (9% of rPeaks), which are likely TSSs of unannotated spliced transcripts. The rPeaks themselves are primarily intergenic, intronic, or antisense TSS-proximal with respect to GENCODE-annotated genes (**Supplemental Fig. S4a, Supplemental Table S3e**). Although this category of rPeaks shows similar levels of evolutionary conservation to the local transcription category of rPeaks, the former category is active in more biosamples and more likely to overlap PacBio TSSs (**Supplemental Fig. S4b-e, Supplemental Table S4**). Additionally, the PacBio reads that overlapped the verified unannotated transcript rPeaks had a similar length distribution to the PacBio reads that overlapped rPeaks in the verified GENCODE TSS category, suggesting that the verified unannotated transcript rPeaks may correspond to lncRNAs missed by GENCODE (**Fig S4f, g)**. To test this hypothesis, we intersected these rPeaks with lncRNA TSSs curated by the lncBook database (Ma et al. 2019) and found that 30% of our verified unannotated transcript rPeaks overlapped the lncBook lncRNA TSSs, a significant enrichment over the local transcription rPeaks and random genomic regions (7% and <1% overlap, respectively, Fisher’s exact test, *p* < 8.8 × 10^-148^, **Supplemental Table S3e**). This result suggests that many of our verified unannotated transcript rPeaks are likely lncRNAs and further expands the growing list of lncRNAs in the human genome.

### RAMPAGE-verified TSSs are enriched for regulatory signatures

We compared our RAMPAGE-verified TSS annotations with GENCODE-annotated TSSs for enrichment in functional, epigenomic, and additional transcriptomic annotations. For these analyses, we only considered genes with both a RAMPAGE-verified unannotated TSS and a GENCODE TSS that did not overlap each other (4,751 genes) and used a uniform 100 bp region centered at an rPeak summit or GENCODE TSS to provide an unbiased comparison.

RAMPAGE-verified unannotated TSSs were more likely to overlap ENCODE candidate cis-regulatory elements (ENCODE Project Consortium et al. 2020) (cCREs, 1.3 fold enrichment, Fisher’s exact test, *p* = 1.7 × 10^-125^) and GTEx expression quantitative trait loci (GTEx Consortium et al. 2017) (eQTLs, 1.2 fold enrichment, Fisher’s exact test, *p* = 2.8 × 10^-6^), compared to matched GENOCDE TSSs (**Fig. 4a**). When we restricted our analysis to RAMPAGE-verified unannotated TSSs active in K562 and their matched GENCODE TSSs, we observed that the verified TSSs were more likely to overlap K562 cCREs (1.8 fold enrichment, Fisher’s exact test, *p* = 8.7 × 10^-92^, **Fig. 4b**) and peaks from the Survey of Regulatory Elements (SuRE) assay, a massively parallel reporter assay testing promoter activity (van Arensbergen et al. 2017) (1.9 fold enrichment, Fisher’s exact tests, *p* = 1.1 x 10^-79^, **Fig. 4a**). The K562 RAMPAGE-verified unannotated TSSs also had higher H3K4me3 and H3K27ac ChIP-seq signals, which had the canonical asymmetric pattern corresponding to transcriptional direction, chromatin accessibility, and Pol II ChIP-seq signals compared to the matched GENCODE TSSs (**Fig. 4c**).

**Figure 4.**
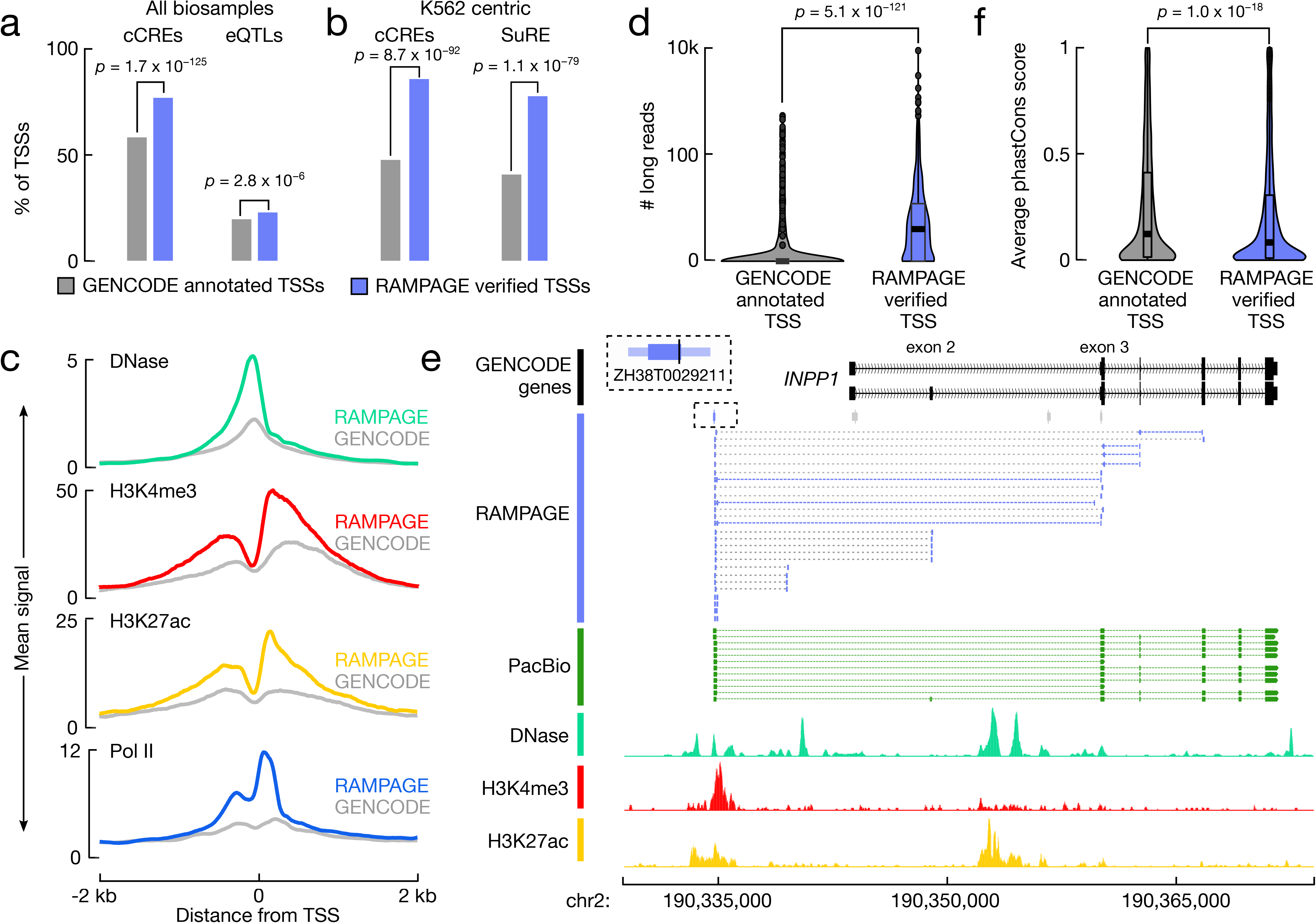
RAMPAGE-verified rPeaks are enriched for regulatory signatures. **a,** Bar plots display the percentage of RAMPAGE-verified TSSs (purple) and matched GENCODE-annotated TSSs (gray) that overlap cell type-agnostic cCREs and the full compendium of GTEx eQTLs. *P*-values are from Fisher’s exact test. **b,** Bar plots display the percentage of RAMPAGE-verified TSSs expressed in K562 (purple) and matching GENCODE-annotated TSSs (gray) that overlap K562 cCREs and SuRE assay peaks. *P*-values are from Fisher’s exact test. **c,** Aggregation plots of epigenomic signals, DNase (teal), H3K4me3 (red), H3K27ac (yellow), and Pol II (blue), at K562 RAMPAGE-verified TSSs (colors) and matched GENCODE-annotated TSSs across a ± 2kb window centered on the summits and TSS, respectively. **d,** Nested violin boxplots showing the number of PacBio 5’ read ends that overlap K562 RAMPAGE-verified TSSs (purple) and matched GENCODE-annotated TSSs (gray). *P*-value is from a Wilcoxon rank sum test. **e,** Genome browser view of *INPP1* locus in K562. RAMPAGE-verified TSS, ZH38T0029211, is linked to the *INPP1* gene by paired-end RAMPAGE reads (purple) whereas the GENCODE-annotated TSSs are not supported by RAMPAGE reads. PacBio reads (green) also support ZH38T0029211 as a verified TSS of *INPP1* and epigenomic signals, DNase (teal), H3K4me3 (red), and H3K27ac (yellow), support promoter activity at ZH38T0029211 and not at the annotated GENCODE TSSs. RAMPAGE rPeaks with RPM > 2 in K562 are shown in purple while those with RPM ≤ 2 are shown in gray. **f,** Nested violin boxplots of average PhastCons conservation scores across RAMPAGE-verified TSSs (purple) and matched GENCODE-annotated TSSs (gray). *P*-value is from a Wilcoxon rank sum test.

We also compared the verified unannotated TSSs with the K562 and GM12878 PacBio long-read RNA-seq data. RAMPAGE-verified unannotated TSSs were more likely to overlap the 5’ ends of PacBio reads compared to the GENCODE-matched controls (median of 3 supporting reads vs. 0 supporting reads, **Fig. 4d**, Wilcoxon test, *p* = 5.1 × 10^-121^). One example is highlighted at the *INPP1* locus (**Fig. 4e**). ZH38T0029211 is a RAMPAGE-verified TSS located 8,844 bp upstream of two GENCODE-annotated TSSs for the *INPP1* gene. The majority of RAMPAGE reads link ZH38T0029211 to the first coding exon (exon 3), while a minority links it to exon 2; similarly, PacBio reads support ZH38T0029211 as a TSS of *INPP1* with the majority also excluding exon 2. Furthermore, epigenomic signals, such as chromatin accessibility and histone ChIP-seq, also support ZH38T0029211 as a novel TSS of *INPP1*.

Despite the enrichment for functional, epigenomic, and transcriptomic annotations, the RAMPAGE-verified unannotated TSSs were less evolutionarily conserved than their matched GENCODE TSSs as measured by phastCons (Siepel et al. 2005) (**Fig. 4f, S4h**; Wilcoxon test, *p* = 2.5 × 10^-19^) and liftOver (Hinrichs et al. 2006) to the mm10 genome (**Supplemental Fig. S4i;** Fisher’s exact test, *p* = 1.5 × 10^-15^). However, the RAMPAGE-verified TSSs were still more conserved than distal enhancer cCREs (cCREs-dELS, **Supplemental Fig. S4h,** Wilcoxon test, *p* = 7.4 × 10^-81^, **Supplemental Fig. S4i**, Fisher’s exact test *p* = 1.2 × 10^-80^) and much more conserved than random genomic regions (Wilcoxon test, *p* < 1.0 × 10^-300^; Fisher’s exact test, *p* < 1.0 × 10^-300^, **Supplemental Fig. S4h, i**). These findings suggest that while the RAMPAGE-verified unannotated TSSs are more biochemically and transcriptionally active in the evaluated cell types, GENCODE TSSs correspond to transcripts expressed in other cell types that have not been surveyed by the RAMPAGE assay. Therefore, for cell type-agnostic data analyses, we suggest users supplement GENCODE TSS annotations with RAMPAGE-annotated TSSs, while for cell type-specific analyses, our results show that RAMPAGE TSSs are a more precise and accurate set of TSSs than the using the entire set of GENCODE annotations.

### RAMPAGE rPeaks identify novel genes that are associated with GWAS phenotypes

We intersected our RAMPAGE rPeaks with variants reported in the NHGRI-EBI genome-wide association study (GWAS) catalog to evaluate the utility of our collection of experimentally derived TSSs (Buniello et al. 2019). Accounting for population-specific linkage disequilibrium (LD, r^2^ > 0.7), our rPeaks overlapped 1,345 variants associated with 208 phenotypes (**Supplemental Table S5a**). To identify disease-associated cell and tissue types, we performed biosample enrichment analysis using our previously published pipeline (ENCODE Project Consortium et al. 2020). However, unlike our previous work, which used nearly one million cCREs, covering -8% of the human genome, our rPeaks had a much smaller genomic footprint; therefore we only observed enrichments passing our FDR thresholds for three phenotypes: (1) obesity related traits; (2) intelligence; and (3) general cognitive ability (see *Methods*; **Supplemental Table S5b**). Generally, enriched cell types were related to disease etiology. For example, intelligence and cognitive ability variants were enriched at rPeaks active in the neuroblastoma cell line SK-N-DZ whereas obesity variants were enriched in rPeaks active in a variety of gastrointestinal and thyroid tissues. This result suggests that while we do not have the power to determine phenotype-relevant cell types for most studies using only RAMPAGE rPeaks, they can still capture biologically relevant enrichments to aid downstream variant interpretation.

Among the 1,345 variants that overlapped RAMPAGE rPeaks, 76% overlapped verified TSSs (50% GENCODE-annotated TSSs and 26% unannotated TSSs) and were therefore linked with an annotated gene by paired-end reads. Of these verified TSS-overlapping variants, 52% were linked with a gene that was not previously reported by the original GWAS and 37% were linked with a gene that was not reported by any GWAS, giving new insights into disease risk (**Supplemental Table S5c**). Of particular interest were RAMPAGE-verified unannotated transcript TSSs that were originally classified as intergenic using GENCODE annotations; these novel TSSs enabled us to assign 41 intergenic SNPs, which were associated with 68 phenotypes, to genes. **Fig. 5a** highlights rs2620666, which is in high LD with two lead SNPs rs750472 and rs13251458 reported to be associated with several cognitive traits (**Supplemental Table S5d**). The original studies reported *FOXH1* and *CYHR1* as possible candidate genes due to their close proximity to the lead SNPs. Although rs2620666 lies only 1,694 bp upstream of a GENCODE-annotated *FOXH1* TSS, it overlaps a RAMPAGE-verified unannotated TSS of *PPP1R16A* (ZH38T0048822, **Fig. 5b**), which encodes a protein phosphatase regulatory subunit. This novel TSS is 11,915 bp upstream of the nearest GENCODE-annotated TSS for *PPP1R16A* and this gene assignment is also supported by PacBio reads (**Fig. 5b, Supplemental Table S5e**). The novel TSS has high RAMPAGE signal in neural cells, brain tissues, and blood cells; moreover, the GTEx consortium identified rs2620666 as an eQTL for several genes (**Supplemental Table S5f**), the most significant of which is *PPP1R16A* in whole blood samples, suggesting that this variant may influence *PPP1R16A* expression. This example highlights the importance of having a comprehensive collection of annotated TSSs so that variants are assigned correctly to the linked genes.

**Figure 5.**
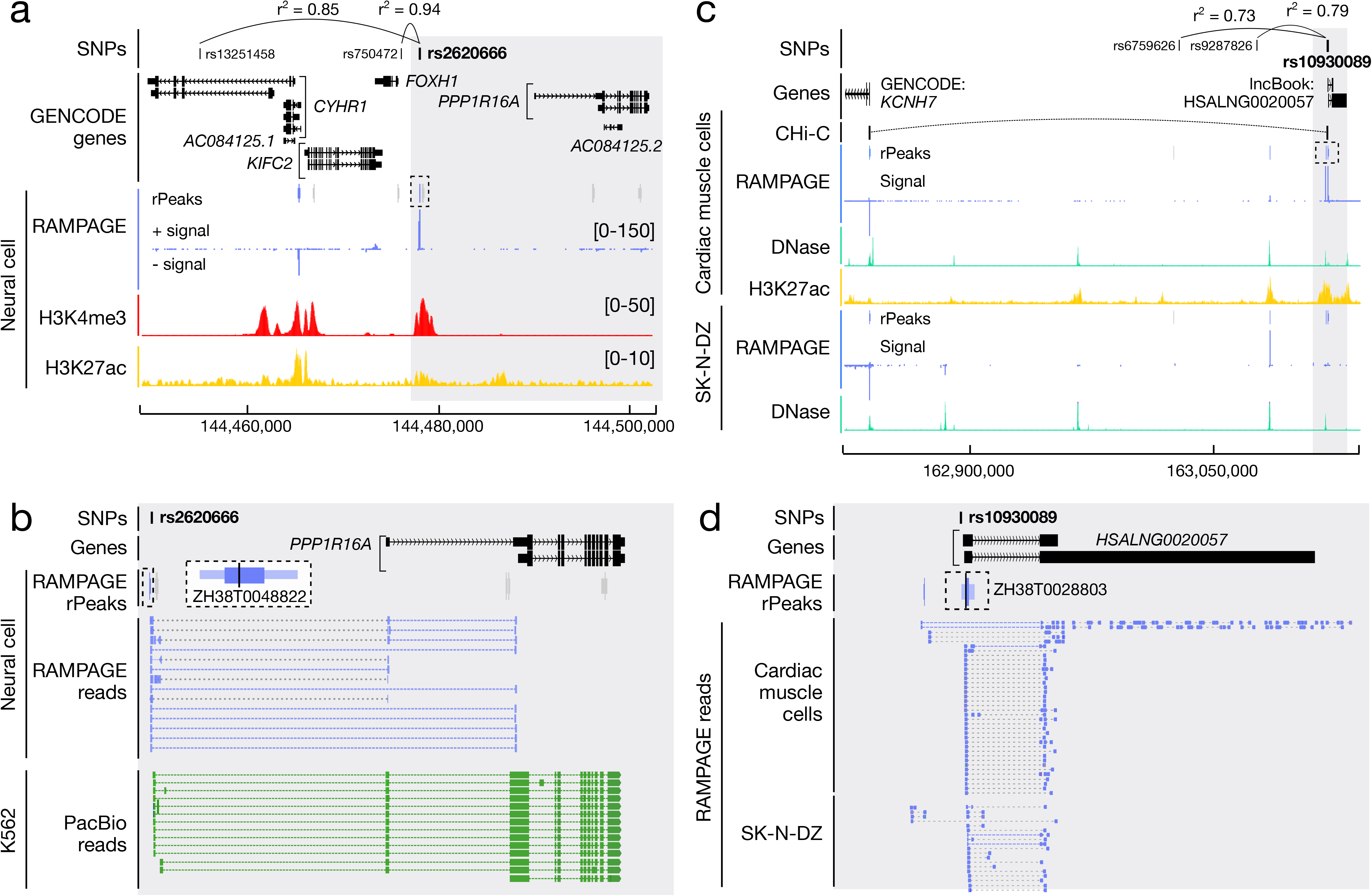
Disease-associated SNPs are linked with new candidate genes using the RAMPAGE rPeak catalog. **a,** Genome browser view of the *CYHR1*-*PPP1R16A* locus. Rs2620666 is in high LD (shown as r^2^ values) with GWAS SNPs rs13251458 and rs750472, and overlaps RAMPAGE rPeak ZH38T0048822 (dashed box). RAMPAGE rPeaks with RPM > 2 in neural cells are shown in purple while those with RPM ≤ 2 are shown in gray; RAMPAGE signal is shown in purple. Supporting epigenomic signals from neural cells, H3K4me and H3K27ac, are shown in red and yellow, respectively. The region shaded in gray is magnified in **b**. **b**, Zoomed-in genome browser view (gray highlight in **a**) displaying RAMPAGE reads (purple) and PacBio reads (green) supporting RAMPAGE peak ZH38T0048822 (dashed box) which is a verified unannotated TSS of *PPP1R16A* and overlaps GWAS SNP rs2620666. RAMPAGE peaks are colored as in **a** and a magnified image of ZH38T0048822 is shown in a larger dashed box with white background. **c,** Genome browser view of the *KCNH7* locus. Rs10930089 is in high LD with GWAS SNPs rs6759626 and rs9287826 and overlaps RAMPAGE rPeak ZH38T0028803 (dashed box). RAMPAGE peaks are colored as described in **a** for cardiac muscle and SK-N-DZ cells. Supporting epigenomic signals from cardiac muscle cells and SK-N-DZ are shown with DNase in teal and H3K27ac in yellow. CHi-C links for cardiac cells are shown in black. The region shaded in gray is magnified in **d**. **d,** Zoomed-in genome browser view (gray highlight in **c**) displaying RAMPAGE reads (purple) from cardiac muscle and SK-N-DZ cells supporting RAMPAGE peak ZH38T0028803 (in dashed box) which overlaps two transcripts of the lncBook lncRNA *HSALNG0020057* and GWAS SNP rs10930089. RAMPAGE peaks shown in purple have RPM > 2 in both cardiac muscle and SK-N-DZ cells.

Finally, we investigated the 34 GWAS variants that overlapped TSSs of RAMPAGE-verified unannotated transcripts (**Supplemental Table S5a**). Of particular interest was rs10930089, an intergenic SNP in high LD with rs6759626 and rs9287826, two lead SNPs associated with general cognitive ability (Davies et al. 2018). Rs10930089 overlaps ZH38T0028803, the TSS of a RAMPAGE-verified unannotated transcript that has high RAMPAGE signal in SK-N-DZ (a neuronal cell line), cardiac tissues, and male reproductive tissues (**Fig. 5c, Supplemental Table S5g**). ZH38T0028803 overlaps the TSSs of two lncRNA transcripts annotated in lncBook, both of which are consistent with the RAMPAGE reads pairs (**Fig. 5d**). In the other direction of the genome, ZH38T0028803 lies 282,766 bp upstream of *KCNH7,* which encodes a potassium voltage channel that has known roles in neurons and the heart (GeneCards Human Gene Database). Variants in *KCNH7* have also been previously associated with bipolar disorder (Strauss et al. 2014) and treatment response in schizophrenia (Wang et al. 2019), suggesting it may play an important role in neuronal pathways. We found that 3D chromatin contact data linked ZH38T0028803 with *KCNH7* in cardiac myocytes (Montefiori et al. 2018) (**Fig. 5c**) but not in iPSC-derived neurons (Rajarajan et al. 2018; Song et al. 2019). Furthermore, ZH38T0028803 has high chromatin accessibility in SK-N-DZ, cardiac cells, and heart tissues, but low chromatin accessibility in fetal brain and iPSC-derived neurons (**Supplemental Table S5h**). Taken together, these results suggest that rs10930089 may modulate the function of ZH38T0028803, the TSS of a lncRNA expressed in neuronal and cardiac cells, and this TSS may also acts as an enhancer for *KCNH7* in both of these two types of cells, with the caveat that the 3D connection is in neuronal cell types other than iPSC-derived neurons.

## Discussion

We annotated 52,546 RAMPAGE rPeaks by integrating 115 RAMPAGE experiments, uniformly curating sites of transcription in hundreds of human cell and tissue types. Using paired-end RAMPAGE reads, we assigned the majority of these rPeaks as TSSs of annotated genes and additionally identified TSSs of over four thousand novel transcripts. We then showed that the TSSs in our catalog were enriched for various regulatory signatures defined using epigenetic and functional data and our catalog complements existing TSS annotations such as those by GENCODE. Through systematic comparisons with CAGE, GRO-cap and PacBio long-read data, we also determined that our catalog of RAMPAGE rPeaks were highly precise and accurate. In particular, PacBio and RAMPAGE had the highest overlap in both GM1878 and K562 cells. PacBio long reads not only supported our RAMPAGE TSS annotations but also supported our assignments of these TSSs to genes (**Fig. 4e, 5b, S3b-d**). PacBio long-read data are particularly advantageous as they allow us to identify novel isoforms and annotate the 3’ ends of transcripts in addition to annotating TSSs. As these data continue to be produced for a wide variety of biosamples by the ENCODE consortium, they will be very useful for further expanding our TSS catalog and enriching transcript annotations.

In both K562 and GM12878 cells, CAGE peaks tended to be the least concordant with the RAMPAGE rPeaks and PacBio 5’ ends (**Supplemental Fig. S2b**). We also noted that CAGE-specific peaks were much more likely to be intronic and intergenic than RAMPAGE rPeaks. However, CAGE peaks were supported by GRO-cap signals at a comparable level as RAMPAGE rPeaks, suggesting that CAGE-specific peaks contain true TSSs (**Supplemental Fig. S2b**). We hypothesize that the CAGE assay can identify a subclass of intergenic and intronic transcription sites, likely eRNAs, that are not detected by RAMPAGE or PacBio long-read RNA-seq. This ability can be used to annotate TSS-distal regulatory elements. Thus, additional comparisons need to be performed with transcription assays that have high rates of eRNA detection, such as BruUV-seq (Magnuson et al. 2015) and PRO-seq/cap (Kwak et al. 2013).

When we compared our catalog of RAMPAGE rPeaks to the FANTOM consortium’s CAGE peak collection, we found that loci missed by RAMPAGE were primarily due to differences in surveyed biosamples (**Fig. 2c**). This result indicates that there is high variability in the transcriptional landscapes among different cell types and a more comprehensive TSS collection can be achieved by surveying a larger collection of biosamples; however, there are additional considerations regarding the composition of a sample collection. Though we currently include over one-hundred biosamples in our RAMPAGE rPeak catalog, the majority of these biosamples are bulk tissue samples which comprise many different cell types. We found that tissue samples generally clustered separately from primary and *in vitro* differentiated cell samples despite some sharing similar biological profiles (**Supplemental Fig. S1f**), possibly due to the technical differences in assaying tissues versus cells. The impact of biosample composition on TSS annotation was also apparent when we observed an enrichment of neuron-related Gene Ontology terms for CAGE-only genes despite the presence of fetal brain tissues and iPSC-derived neurons in our RAMPAGE sample collection. This result suggests that these early developmental brain tissues may be dominated by precursor cells such as immature neuronal progenitors or radial glia and that the iPSC-derived neurons may represent alternative cell states from mature neurons. Interestingly, on a genome-wide scale, SK-N-DZ has a transcriptional profile that is more similar to iPSC-derived neurons than to mature neurons, as evident from UMAP embedding (**Supplemental Fig. S1f**). The discrepancy among the different types of neuronal cells was further highlighted by our GWAS analysis where we observed that the cognitive phenotype related SNPs overlapped a novel TSS active in SK-N-DZ cells but not in iPSC-derived neurons. Therefore, although SK-N-DZ cells overall share similar transcriptomic signatures to iPSC-derived neurons, there are subtle differences in cellular state that may have important impacts on variant and disease interpretation. With further developments of single-cell transcriptomic technologies to capture the 5’end of transcripts, it will be important to expand our TSS identification methods to build a comprehensive catalog by cell type, particularly in heterogeneous tissues such as the brain.

Even though we observed enrichments in some tissues for GWAS variants associated with three phenotypes, our comparisons were underpowered compared to our previous work (ENCODE Project Consortium et al. 2020) due to the small genomic footprint of RAMPAGE rPeaks. Despite this, we demonstrated that accurate TSS annotations, particularly those TSSs linked with known transcripts, are important for interpreting variants reported by GWAS. Additionally, we anticipate that such collections will also be important for the detection and interpretation of rare and *de novo* variants uncovered by whole-genome sequencing efforts, as these variants have larger effect sizes and may be more likely to fall within promoter regions than in distal regulatory elements. For example, a recent study found an enrichment of *de novo* variants associated with autism spectrum disorder in promoters (An et al. 2018). Therefore, accurate, cell type-specific TSS annotations can improve our power for interpreting the impact of *de novo* genetic variation across cell types.

Finally, we identified 4,129 TSSs for unannotated transcripts, many of which we hypothesize to be lncRNAs although we could not test this hypothesis with only the beginning portion of these transcripts. It is also unclear if these transcripts carry out any cellular functions. A wide range of functional mechanisms have been reported for lncRNAs, varying from transcriptional regulation of other genes via epigenetic or antisense means to simply being the byproducts of strong enhancers (Quinn and Chang 2016; Fang and Fullwood 2016). With development of antisense oligonucleotide (ASO) and CRISPR perturbation technologies, it is now possible to perform screens to identify functional lncRNAs in a high-throughput manner (Joung et al. 2017; Liu et al. 2017; Ramilowski et al. 2020). As these collections of functionally validated lncRNAs become available across diverse cellular contexts, we plan to further refine our TSS catalog to include such functional information.

In summary, our catalog of RAMPAGE rPeaks expands the human transcriptional landscape across over one hundred cell and tissue types. The catalog provides a valuable resource to the biological community by improving annotations for studying gene regulation and aiding in the interpretation of genetic variants associated with human diseases.

## Acknowledgements

We thank Gabriela Balderrama-Gutierrez, Diane Trout, and Julien Lagarde for discussions on how to best analyze TSSs from long read PacBio data. This work was supported by grants from the NIH under U24HG009446 and UM1HG009443.

## Supplemental Tables

Supplemental Table S1. Summary of RAMPAGE datasets and rPeaks

Supplemental Table S2. Gene ontology enrichment for CAGE-only and RAMPAGE-only genes

Supplemental Table S3. Assignment of RAMPAGE rPeaks to genes

Supplemental Table S4. Enrichment for RAMPAGE rPeaks features stratified by TSS class

Supplemental Table S5. Overlap of GWAS variants with RAMPAGE rPeaks

## Methods

### Generating a collection of RAMPAGE rPeaks

#### Curating RAMPAGE experiments

As of September 1, 2020 there were 155 ENCODE3 RAMPAGE experiments at the ENCODE portal (https://www.encodeproject.org/search/?type=Experiment&status=released&perturbed=false&assay_title=RAMPAGE&award.rfa=ENCODE3&perturbed=true). From the portal, we downloaded RAMPAGE bam alignment files, which contained reads mapped to the GRCh38/hg38 reference genome by the ENCODE Data Coordination Center using STAR (https://www.encodeproject.org/data-standards/rampage/). We then removed redundant reads as described in Zhang et al. (Zhang et al. 2019), briefly summarized as follows. First, properly aligned read pairs (R1 and R2 denote each mate of a read pair) with uniquely aligned R2 reads were collapsed with the same alignment coordinates and the identical 15-bp barcode at the 5’-end of R2 reads to remove PCR duplicates. We then pooled read pairs from biological replicates together after the PCR duplicate removal and created signal bigWig files of the 5’ ends of R1 reads that we used for all subsequent signal quantifications (available for download on our companion site). Finally, we excluded all experiments with a non-redundancy fraction less than 0.25, which resulted in a final collection of 115 high quality RAMPAGE experiments (Supplemental Table S1).

Relevant script: rm_pcr.py

#### Calling RAMPAGE peaks

We called RAMPAGE peaks as described in Zhang *et al. (Zhang et al. 2019)*. Briefly, RAMPAGE peaks were clustered with the 5’-most base of aligned R1 reads using F-seq (Boyle et al. 2008) (parameter: feature length = 30 and fragment size = 0). For each peak, we identified a high-density region, which contained 70% of the reads in each original peak, and a summit, which was the genomic position with the highest number of R1 5’ read ends.

Relevant script: call_peak.py

#### Filtering RAMPAGE peaks

For each RAMPAGE experiment, the Gingeras lab also performed a matching total RNA-seq experiment on the same biosample, which we used to filter RAMPAGE peaks. Using bigWigAverageOverBed, we calculated the total RNA-seq and RAMPAGE signals (column four of the resulting file, sum) across each RAMPAGE peak. We excluded peaks whose RNA-seq signals were greater than their RAMPAGE signals (i.e., peaks that fell below the x=y line, **Supplemental Fig. S1**). These peaks predominantly overlapped annotated exons. Finally, to further select for high-quality annotations, we only retained peaks with RPM (reads per million) > 2, which resulted in a set of 1,147,456 peaks across all experiments with an average of 9,978 per experiment (**Supplemental Table S1**).

Relevant script: 1_Peak-Filtering.sh

#### Generating representative RAMPAGE peaks

To generate representative RAMPAGE peaks (RAMPAGE rPeaks), we adapted the representative DNase Hypersensitivity Site (rDHS) pipeline as described by the ENCODE Project Consortium (ENCODE Project Consortium et al. 2020). First, to retain strand-specific information, we separated peaks based on DNA strand, and then clustered the strand-specific peaks across all 115 experiments using *bedtools merge (Quinlan and Hall 2010)*. For each cluster, we selected the peak with the highest RPKM (reads per kilobase per million) signal as the rPeak. All peaks that overlapped this rPeak-as defined by using *bedtools intersect* with default parameters-were then removed. We iteratively repeated this process until all 1.1M RAMPAGE peaks were represented by a collection of 80,157 non-overlapping rPeaks. To reduce false positives, we discarded all singleton rPeaks (i.e., rPeaks that represented only one experiment) unless they had an RPM > 5, resulting in a final set of 52,546 rPeaks.

Relevant script: 2_rPeak-Annotation.sh

### Genomic context and enrichment

#### Determining genomic context

We used the following hierarchical approach to assign genomic contexts to annotations (including RAMPAGE rPeaks and FANTOM CAGE peaks). We used *bedtools intersect* to determine overlapping features with overlap requirements as described below:

1. *TSS-overlapping*: rPeak overlapped an annotated TSS from GENCODEv31 basic annotations. Use default parameters for *bedtools intersect*.
2. *TSS-Proximal*: rPeak fell within ± 500 bp of an annotated TSS from GENCODEv31 basic. Required at least 50% of RAMPAGE rPeak to overlap region (-f 0.5).
3. *Exon*: rPeak overlapped “exon” annotation from GENCODEv31 basic which include coding exons (CDS), exons of non-coding genes, and untranslated regions (UTRs). Required at least 50% of RAMPAGE rPeak to overlap exon (-f 0.5).
4. *Intron:* rPeak overlapped an annotated gene from GENCODEv31 basic but not an exon. Required at least 50% of RAMPAGE rPeak to overlap gene (-f 0.5).
5. *Intergenic:* all remaining rPeaks

Relevant script: Determine-Genomic-Context.sh

#### Assigning strand

1. *TSS-overlapping*: assign strand of the transcript the RAMPAGE rPeak overlaps. If the RAMPAGE rPeak overlaps TSSs on both strands, the strand matching the rPeak is assigned.
2. *Proximal*: assign strand of the transcript RAMPAGE rPeak falling within 500 bp. If the RAMPAGE rPeak overlaps TSSs on both strands, the strand matching the rPeak is assigned.
3. *Exon*: assign strand of the transcript containing the exon the rPeak overlaps. If the RAMPAGE rPeak overlaps exons on both strands, the strand matching the rPeak is assigned.
4. *Intron:* assign strand of the transcript the rPeak overlaps. If the RAMPAGE rPeak overlaps transcripts on both strands, the strand matching the rPeak is assigned.
5. *Intergenic:* assign strand of the closest transcript as determined by *bedtools closest* using the TSS basic annotations. If the RAMPAGE rPeak is equally close to transcripts on both strands, the strand matching the rPeak is assigned.

Relevant script: Determine-Genomic-Context.sh

#### Determining genomic context enrichment

To determine the genomic background, we calculated the percentage of the GRCh38 genome comprising each of the annotations: (TSS: 0.0004%; TSS-proximal: 2.2%; Exon: 3.7%; Intron: 45.5%; Intergenic: 48.6%). We then determined the percentage of total rPeaks falling in each annotation and calculated fold enrichment.

Relevant script: Determine-Genomic-Context.sh

### Boundary and summit analysis

For each rPeak, we calculated the median peak boundary, high-density boundary and summit variation for each peak that was represented. We did not include peaks that were selected as the rPeaks in this analysis.

Relevant script: Compare-Boundary-Variation.sh

### UMAP

We performed two separate UMAP analyses: one using all 115 biosamples (**Figure S1f**) and one using the subset of all 87 tissue samples (Figure 1g). For each biosample, we calculated the RPKM (reads per kilobase per million) at each rPeak. We then combined these results to create two input matrices, 52,547 by 115 and 52,547 by 87, respectively, where each row is a RAMPAGE rPeak and each column is a biosample. For each entry of the matrix we took the log10 of each entry and normalized each row using sklearnStandardScaler. We then implemented the UMAP algorithm using the python UMAP-learn package with n_neighbors = 10 and default values for the remaining parameters.

Relevant script: UMAP-Analysis.sh

### Comparisons with other transcription annotations

#### Comparison with CAGE peaks

We downloaded CAGE peaks and quantifications from the FANTOM consortium:

- Peaks: https://fantom.gsc.riken.jp/5/datafiles/reprocessed/hg38_latest/extra/CAGE_peaks/hg38_fair+new_CAGE_peaks_phase1and2.bed.gz
- Quantifications: https://fantom.gsc.riken.jp/5/datafiles/reprocessed/hg38_latest/extra/CAGE_peaks_expression/hg38_fair+new_CAGE_peaks_phase1and2_tpm.osc.txt.gz

To compare the overall concordance of peak collections, we intersected the entire collection of CAGE peaks with the entire collection of RAMPAGE peaks using *bedtools intersect* with the requirement that at least 25% of the CAGE peak overlapped the RAMPAGE peak and the peaks fell on the same strand.

Relevant script: RAMPAGE-CAGE-All-Peak-Comparison.sh

To extract peaks active in K562 and GM12878, we selected all peaks with an average TPM (transcripts per million) > 2 across the three surveyed replicates (columns 563-565 for K562 and columns 171-173 for GM12878). We compared these peaks with RAMPAGE rPeaks with RPM > 2 in K562 and GM12878, respectively, using *bedtools intersect*, requiring overlapping peaks to be on the same strand and overlap a minimum of 25% of the CAGE peak.

Relevant script: Extract-CAGE-Peaks.sh

#### Comparison with PacBio long-read RNA-seq data

We downloaded the following bam files from the ENCODE project data portal: ENCFF546DOT and ENCFF709YES for K562 and ENCFF247TLH, ENCFF431IOE, ENCFF520MMC, and ENCFF626GWM for GM12878. We merged and sorted bam files for each cell type, split reads by genomic strand, and used *bedtools bamtobed* to extract the 5’ ends of reads.

Relevant script: Format-PacBio-Data.sh

We used *bedtools intersect* with default parameters to intersect PacBio 5’ read ends with RAMPAGE and CAGE peaks. To only count strand matching intersections, RAMPAGE and CAGE peaks were first split by strand and then intersected with 5’ ends on the same strand.

Relevant script: Compare-TSS-Annotations.sh

#### Comparison with GRO-cap signal

We downloaded the following GRO-cap signal files from GEO under accessions GSM1480321 and GSM1480323 for K562 and GM12878, respectively:

- GSM1480321_K562_GROcap_wTAP_plus.bigWig
- GSM1480321_K562_GROcap_wTAP_minus.bigWig
- GSM1480323_GM12878_GROcap_wTAP_plus.bigWig
- GSM1480323_GM12878_GROcap_wTAP_minus.bigWig

To calculate average signal at RAMPAGE rPeaks, CAGE peaks, and PacBio 5’ ends, we lifted down the 1 bp summits or read ends to the hg19 genome using the UCSC liftOver tool (Kuhn et al. 2013) with default parameters. We then set region width to a uniform 50 bp centered on the peak summits or 5’ ends and, using the UCSC bigWigAverageOverBed function, calculated the average signal across each region.

To determine a signal threshold for high GRO-cap signal, we first randomly selected 500k 50 bp genomic regions and calculated their average GRO-cap signal. We then selected the 99.5th percentile as the threshold for *high signal* which was 0.06 in K562 and 0.08 in GM12878, respectively.

Relevant script: Calculate-GROcap-Signal.sh

#### Comparison of GENCODE covered genes

We first set peak width to a uniform 100 bp centered around each peak summit or 5’ read end and then intersected these regions with annotated TSSs of GENCODE 31 genes using default *bedtools intersect* and requiring annotations to be on the same strand (-s flag). We performed gene ontology analysis using PantherDB’s online database (Mi et al. 2017). We first performed this analysis for the entire sets of RAMPAGE and CAGE peaks, then for peaks and PacBio 5’ read ends in K562 and GM12878 cells.

Relevant scripts: RAMPAGE-CAGE-Gene-Comparison.sh,

RAMPAGE-CAGE-PacBio-Gene-Comparison.sh

### Assigning RAMPAGE rPeaks to Genes

#### Curating verified GENCODE TSSs, verified unannotated TSSs, unannotated transcript TSSs and local transcription rPeaks

We developed the following computational workflow to link RAMPAGE rPeaks with genes, which is detailed in **Figure S3a**. Briefly, based on the genetic context of the rPeak and the location of its supporting 3’ reads, we assigned the rPeak into one of six categories:

1. *Verified GENCODE TSS*: rPeak overlaps an annotated GENCODE TSS and its 3’ read ends overlap a downstream exon.
2. *Verified unannotated TSS*: rPeak does not overlap an annotated GENCODE TSS (i.e., rPeak is either TSS-proximal, exonic, intronic, or intergenic) and its 3’ read ends overlap a downstream exon.
3. *Candidate GENCODE TSS*: rPeak overlaps a TSS, first exon or is TSS-proximal to either a single exon transcript, or a transcript with a first exon greater than 500 nt.
4. *Unannotated transcript TSS*: rPeak is supported by reads with 3’ ends that do not overlap an annotated GENCODE exon.
5. *Local transcription*: rPeak is supported by reads that span less than 1 kb or map to the first exon of the transcript.
6. *Discard*: We discarded all rPeaks that overlapped exons that were not the first exon of a transcript or only supported by reads that spanned more than 500 kb.

Relevant scripts: Analyze-RAMPAGE-Read-Mates.sh

Assign-Gene-Links.sh

#### Overlap of novel transcripts with lncRNAs

We downloaded lncRNA annotations (lncRNA_LncBook_GRCh38_9.28.gtf) from lncBook (Ma et al. 2019) (https://bigd.big.ac.cn/lncbook/index) and extracted annotated TSSs. Then, we intersected RAMPAGE rPeaks using default *bedtools intersect* parameters and requiring annotations to be on the same strand (-s flag). We also calculated the overlap of lncBook TSSs with 500k 100 bp random genomic regions generated using *bedtools random*.

Relevant script: Intersect-lncBook-TSSs.sh

### Comparison of GENCODE and verified TSSs

#### Generating sets of matched GENCODE TSSs

We first selected all GENCODE genes that did not have a single annotated TSS overlapping a RAMPAGE rPeak. Of these, we then selected all genes with a RAMPAGE-verified TSS. Because of the no overlapping requirement, these RAMPAGE-verified TSSs were either TSS-proximal, exonic, intronic, or intergenic. The GENCODE-annotated TSSs of these genes served as the matched GENCODE TSS set. In total, we curated 8,391 GENCODE-matched TSSs to compare with 6,243 RAMPAGE-verified TSSs. We also curated K562-specific annotations by selecting all RAMPAGE-verified TSSs with an RPM > 2 in K562 and their matched GENCODE TSSs, resulting in a set of 1,768 GENCODE-matched TSSs to compare with 966 RAMPAGE-verified TSSs in K562 cells.

Unlike the RAMPAGE-verified TSSs, GENCODE TSSs were only 1 bp in width; therefore, to eliminate biases due to region width, we generated uniform 100 bp regions centered on either RAMPAGE -verified TSS summits or GENCODE TSSs, respectively.

Relevant script: Compare-Verified-TSS-GENCODE.sh

#### Overlap of RAMPAGE-verified and matched GENCODE TSSs with ENCODE cCREs

We downloaded cell type-agnostic cCREs and K562-specific cCREs from the ENCODE SCREEN database (screen.encodeproject.org). For the K562 cCREs, we filtered out “Low-DNase” cCREs, which are regulatory regions deemed inactive in the cell type. Using *bedtools intersect* with default parameters and the -u flag to count unique elements, we intersected the uniform 100 bp sized TSS regions (as described above) with the cell type-agnostic cCREs. We repeated this analysis using the K562 cCREs and the uniform 100 bp sized K562 regions.

Relevant script: Compare-Verified-TSS-GENCODE.sh

#### Overlap of RAMPAGE-verified and matched GENCODE TSSs with GTEx eQTLs

We downloaded eQTLs from the GTEx database (GTEx_Analysis_v8_eQTL.tar), aggregated across all *signif_variant_gene_pairs.txt files, and reformatted the results into bed format. Using *bedtools intersect* with default parameters and the -u flag to count unique elements, we intersected the uniform 100 bp sized TSS regions (as described above) with the eQTL bed file.

Relevant script: Compare-Verified-TSS-GENCODE.sh

#### Overlap of K562 RAMPAGE-verified and matched GENCODE TSSs with SuRE peaks

We downloaded SuRE peaks from the Supplementary Data section of van Arensbergen *et al. (van Arensbergen et al. 2017)* (SuRE-peaks_K562.45.55_raw_sep_globalLambda.annotated_LP160616.txt). We reformatted this file into bed format, and lifted the regions from the hg19 genome up to the hg38 genome using UCSC’s liftOver tool with default parameters and the hg19ToHg38.over.chain chain file. We then intersected the uniform 100 bp sized TSS regions from K562 (as described above) with hg38 K562 SuRE peaks using *bedtools intersect* with default parameters and the -u flag to count the number of unique regions that overlapped.

Relevant script: Compare-Verified-TSS-GENCODE.sh

#### Aggregate epigenomic signals at RAMPAGE-verified and matched GENCODE TSSs

Using 1 bp bins, we calculated the average DNase-seq, and H3K4me3, H3K27ac and Pol II ChIP-seq signals, along a 4 kb window centered across the RAMPAGE-verified rPeak summit or matched GENCODE TSS, respectively, accounting for strand orientation. We used the following uniformly processed bigWig files from the ENCODE portal:

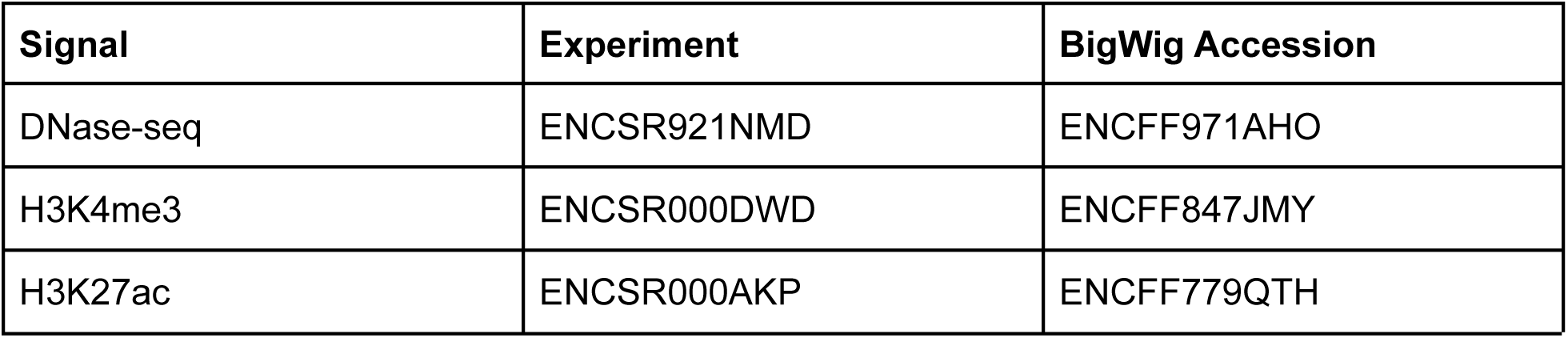
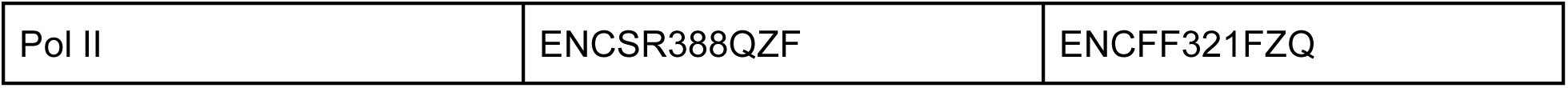

| Signal | Experiment | BigWig Accession |
| --- | --- | --- |
| DNase-seq | ENCSR921NMD | ENCFF971AHO |
| H3K4me3 | ENCSR000DWD | ENCFF847JMY |
| H3K27ac | ENCSR000AKP | ENCFF779QTH |
| Pol II | ENCSR388QZF | ENCFF321FZQ |

Relevant script: Run-Aggregate-Signal.sh

#### Overlap of RAMPAGE-verified and matched GENCODE TSSs with PacBio 5’ read ends

We intersected the uniform 100 bp sized TSS regions from K562 (as described above) with K562 PacBio 5’ read ends (generated from Format-PacBio-Data.sh, see above for more details) using *bedtools intersect* with the -c flag-which counts the number of overlapping entries-and requiring genomic strands to match.

Relevant script: Compare-Verified-TSS-GENCODE.sh

#### Conservation of RAMPAGE-verified and match GENCODE TSSs

We calculated the average PhastCons conservation (100way vertebrate) across the uniform 100 bp regions (as described above) using UCSC’s bigWigAverageOverBed (Kent et al. 2010).

We lifted the uniform 100 bp sized TSS regions (as described above) over to the mm10 genome using UCSC’s liftOver tool (Hinrichs et al. 2006) with a minMatch = 0.5 and the hg38ToMm10.over.chain chain file. We then calculated the percentage of total regions that successfully lifted over. We also compared the liftOver rates of ENCODE cCREs-dELS-extracted from the cell type-agnostic set of cCREs-and 500k random regions of the genome generated from *bedtools random*. For comparison, both these sets of regions were resized to 100 bp around the region center.

Relevant script: Compare-Verified-TSS-GENCODE.sh

### Interpreting GWAS variants with the RAMPAGE rPeak catalog

#### Overlap of GWAS variants

We curated SNPs reported by the NHGRI-EBI GWAS catalog as of January 2019 and using population specific linkage disequilibrium, incorporating all SNPs in high LD (r^2^ > 0.7) with this collection, as described in (ENCODE Project Consortium et al. 2020). We created a master bed file with these annotations and intersected them with our RAMPAGE rPeak catalog using default bedtools parameters. To compare gene assignments, we extracted reported and mapped genes from the original studies (columns 14 and 15 from the downloaded NHGRI-EBI GWAS catalog file) and determined if our rPeak linked genes (from read pair analysis) were represented on the list.

Relevant script: Overlap-GWAS-SNPs.sh

#### Cell type enrichment

We tested whether sets of GWAS SNPs were enriched in RAMPAGE rPeaks activity in specific biosamples using the same GWAS enrichment pipeline as described in (ENCODE Project Consortium et al. 2020). Because RAMPAGE rPeaks have a much smaller genomic footprint than other collections of genomic regions (e.g., cCREs), we only included studies for which at least 15 LD blocks contained a SNP that overlapped a RAMPAGE rPeak (67 out of 397 initially tested GWAS studies). We reported all enrichments with an FDR corrected *p*-value less than 0.5 (**Supplemental Table S5b**).

Relevant repository: https://github.com/weng-lab/VIPER

#### 3D chromatin interactions between ZH38T0028803 and KCNH7

We downloaded the cardiomyocyte promoter capture Hi-C data (Montefiori et al. 2018) from ArrayExpression under the accession E-MTAB-6014:

• E-MTAB-6014.processed.1.zip -> capt-CM-replicated-interactions-1kb.bedpe and iPSC neuron promoter capture Hi-C data (Song et al. 2019) from GEO under the accession GSE113481:

• GSM3106832_cortical.cutoff.5.washU.txt.gz

• GSM3598046_hippocampal.cutoff.5.washU.txt.gz

• GSM3598048_motor.cutoff.5.washU.txt.gz

We also requested iPSC neuron Hi-C loop calls directly from Rajarajan *et al*. (Rajarajan et al. 2018), who generously provided these annotations.

Using *bedtools intersect* with default parameters, we intersected links with the *KCNH7* locus, requiring one of the *KCNH7* GENCODE TSSs to overlap one anchor and ZH38T0028803 to overlap the other anchor. Because the cardiomyocyte data was mapped to the hg19 genome, we lifted down the *KCNH7* TSSs and ZH38T0028803 coordinates to hg19 using UCSC liftOver with default parameters.

Relevant script: Compare-3D-Chromatin-Links.sh

---

**Supplemental Figure S1.**
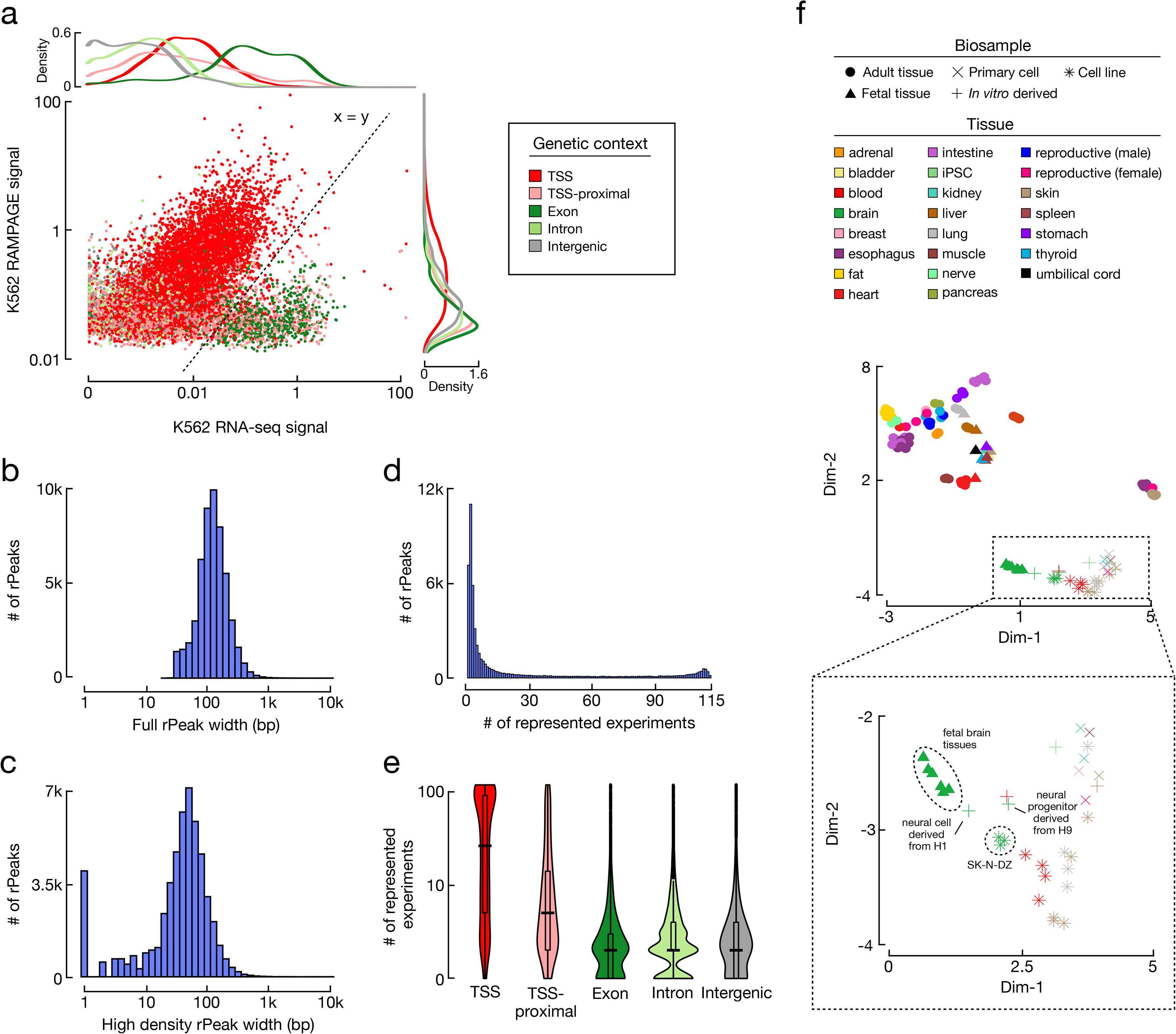
General properties of RAMPAGE rPeaks. **a,** A scatterplot comparing the RNA-seq signals (x-axis) and RAMPAGE signals (y-axis) in K562 cells at RAMPAGE peaks identified in K562. Points are colored by genomic context: TSS in red, TSS-proximal in pink, exon in dark green, intron in light green and intergenic in grey. Density plots along the x- and y-axes show the distributions of signal stratified by genomic context for RNA-seq and RAMPAGE signals, respectively. The dashed line represents the x=y line used to filter RAMPAGE peaks prior to the rPeak pipeline; peaks falling below this line were excluded. **b,** A histogram depicting the distribution of rPeak full-peak widths. **c,** A histogram depicting the distribution of rPeak high-density region widths. **d,** A histogram depicting the number of RAMPAGE experiments represented by each rPeak. **e,** A violin-boxplot depicting the number of experiments represented by each rPeak stratified by genomic context as in **a**. **f**, Scatterplot displaying a two-dimensional Uniform Manifold Approximation and Projection (UMAP) embedding of 115 biosamples using RAMPAGE signal across all rPeaks as input features. Markers are shaped by biosample category and colored by tissue of origin as defined in the legend.

**Supplemental Figure S2.**
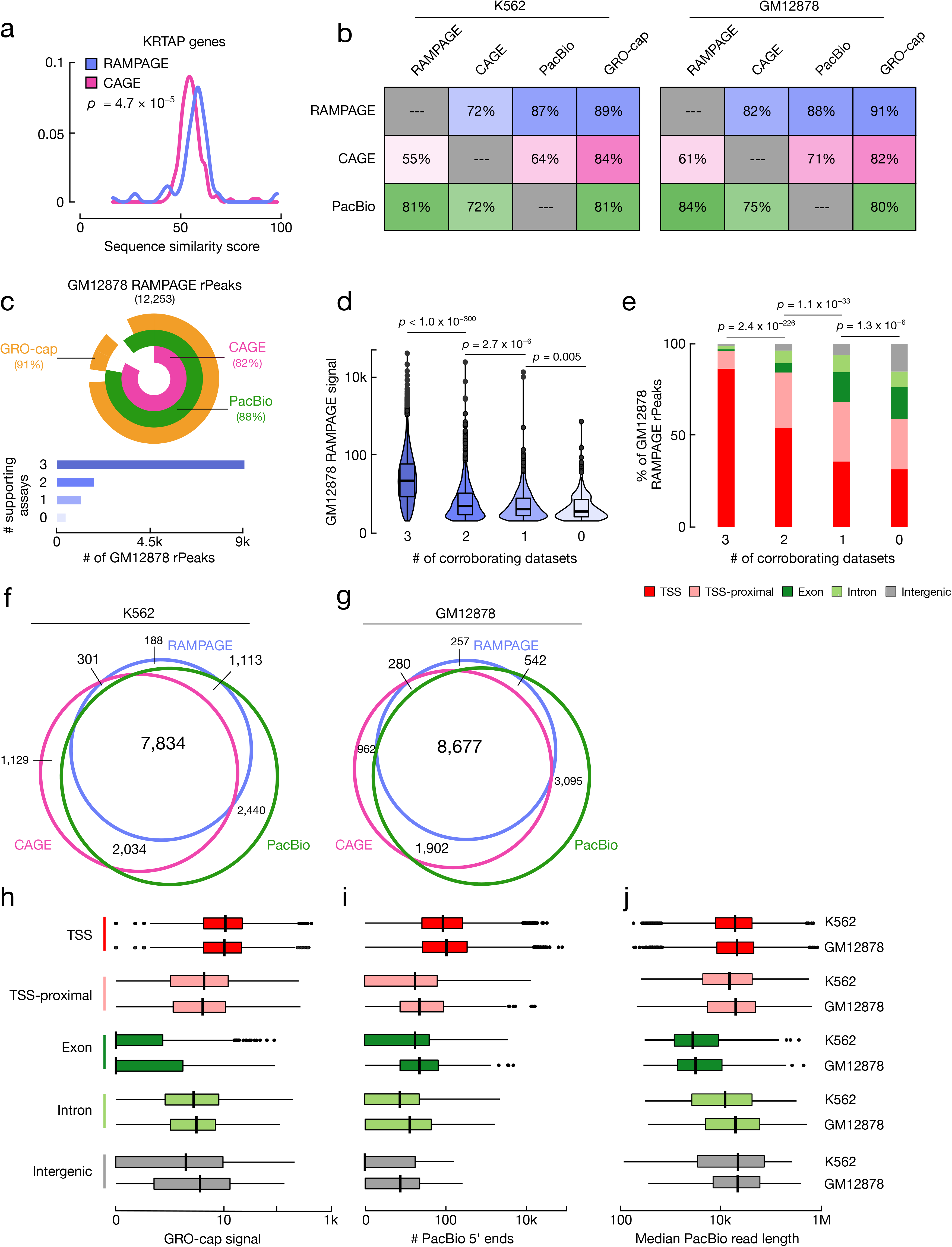
Comparison of RAMPAGE rPeaks with individual transcriptome annotations. **a,** A density plot showing the distributions of the similarity scores for sequences surrounding the TSSs of RAMPAGE-only (purple) and CAGE-only (pink) KRTAP genes. Sequence similarity is calculated by taking the maximum score of all pairwise local alignments. *P-*value corresponds to a two-sided Wilcoxon test. **b,** A heatmap displaying the percentage of RAMPAGE rPeaks (purple), CAGE peaks (pink) and PacBio 5’ read ends (green) that overlap each other and have high GRO-cap signal in K562 (left) and GM12878 (right) cells. **c,** (top) VennPie diagram displaying the percentage of GM12878 RAMPAGE rPeaks that overlap K562 CAGE peaks (pink) or PacBio 5’ ends (green), or have high GRO-seq signals (orange). Concentric circles show the percentages that are similarly supported between the three assays. (bottom) Bar plot with the number of GM12878 rPeaks stratified by the number of supporting transcriptomic assays as described in the above VennPie. **d,** Violin-boxplot showing the distributions of the average GM12878 RAMPAGE signal across rPeaks stratified by the number of supporting assays as defined in **c.** *P*-values correspond to two-sided pairwise Wilcoxon tests with FDR correction. **e,** Stacked bar graphs showing the percentage of GM12878 rPeaks belonging to each genomic context (TSS: red, TSS-proximal: pink, exon: dark green, intron: light green, intergenic: gray) stratified by the number of supporting assays as defined in **c.** *P*-values correspond to Chi-square tests. **f,** Venn diagram showing the overlap of genes with TSSs that overlap either RAMPAGE rPeaks (purple), CAGE peaks (pink) or PacBio 5’ ends (green) in K562. **g,** Venn diagram as described in **f** for annotations in GM12878. **h**, Boxplots showing the GRO-cap signal at RAMPAGE rPeaks stratified by genomic context in K562 (top) and GM12878 cells (bottom). **i**, Boxplots plots showing the number of PacBio 5’ ends that overlap RAMPAGE rPeaks stratified by genomic context in K562 (top) and GM12878 cells (bottom). **j**, Boxplots plots showing the median lengths of the PacBio reads with 5’ ends that overlap each RAMPAGE rPeak stratified by genomic context in GM12878 (top) and K562 cells (bottom).

**Supplemental Figure S3.**
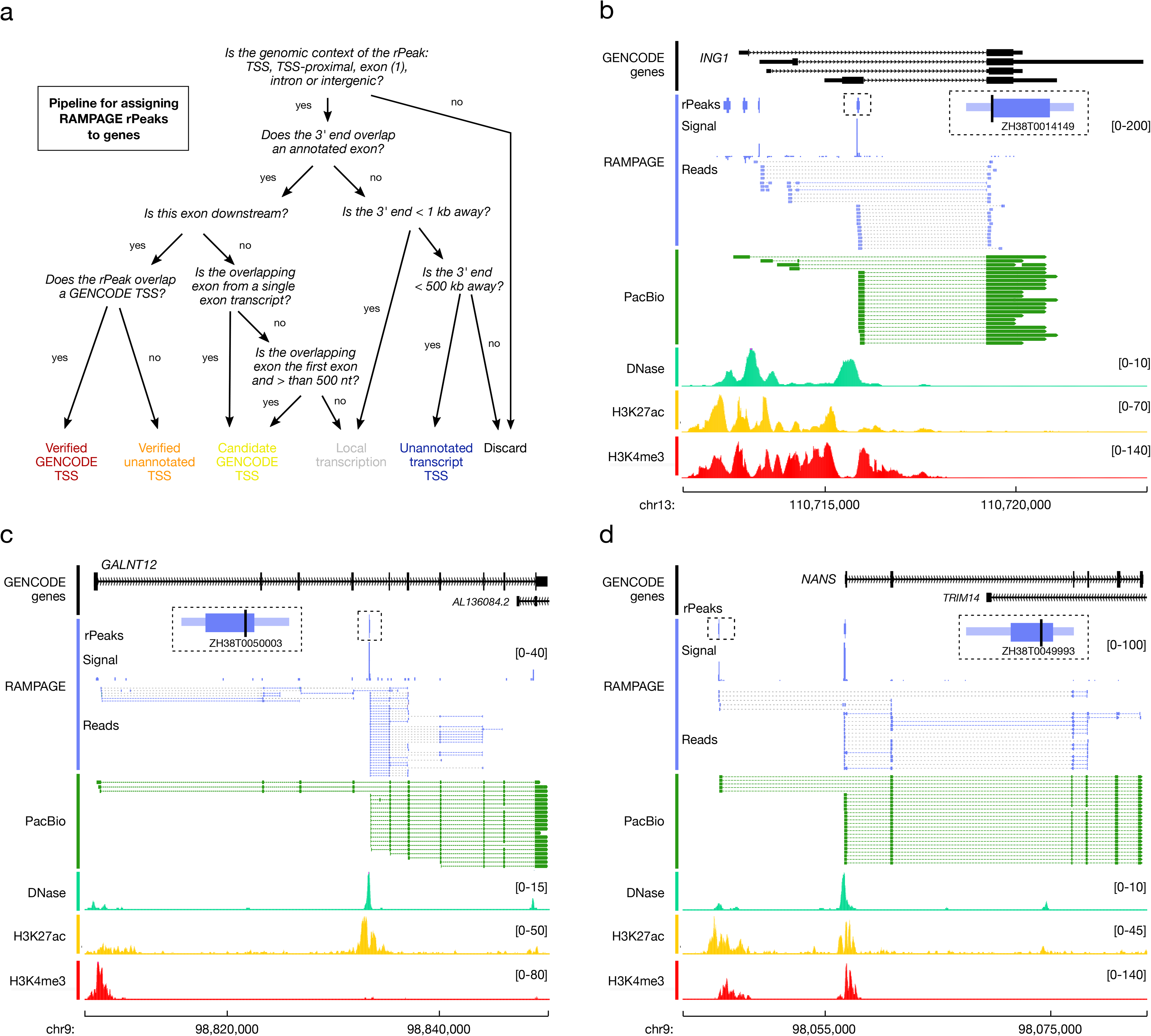
Examples of exonic, intronic, and intergenic rPeaks in K562 cells. **a,** Decision tree depicting the computational workflow for assigning RAMPAGE rPeaks to genes. **b,** Genome browser view of the *ING1* locus. Exonic RAMPAGE rPeak ZH38T0014149 (in a small dashed box, with a magnified version shown to its right) overlaps an annotated exon of *ING1*, but is also a novel TSS for this gene. This annotation is supported not only by RAMPAGE signal and reads (purple), but PacBio reads (green) and epigenomic signals (DNase: teal; H3K27ac: yellow; H3K4me3: red) in K562. Read pairs are denoted by dashed gray lines and split reads (i.e., single reads that span splice junctions) are denoted by colored lines (purple for RAMPAGE and green for PacBio). **c,** Genome browser view of the *GALNT12* locus. Intronic RAMPAGE rPeak ZH38T0050003 (the smaller dashed box, with a magnified version shown to its left), overlaps an annotated intron of *GALNT12*, but is also a novel TSS for this gene. This annotation is supported not only by RAMPAGE signal and reads (purple), but PacBio reads (green) and epigenomic signals (colored as in **b**) in K562. **d,** Genome browser view of the *NANS* locus. Intergenic RAMPAGE rPeak ZH38T0049993 (the smaller dashed box, with a magnified version shown to the right), lies upstream of *NANS*, but is also a novel TSS for this gene. This annotation is supported not only by RAMPAGE signal and reads (purple), but PacBio reads (green) and epigenomic signals (colored as in **a**) in K562.

**Supplemental Figure S4.**
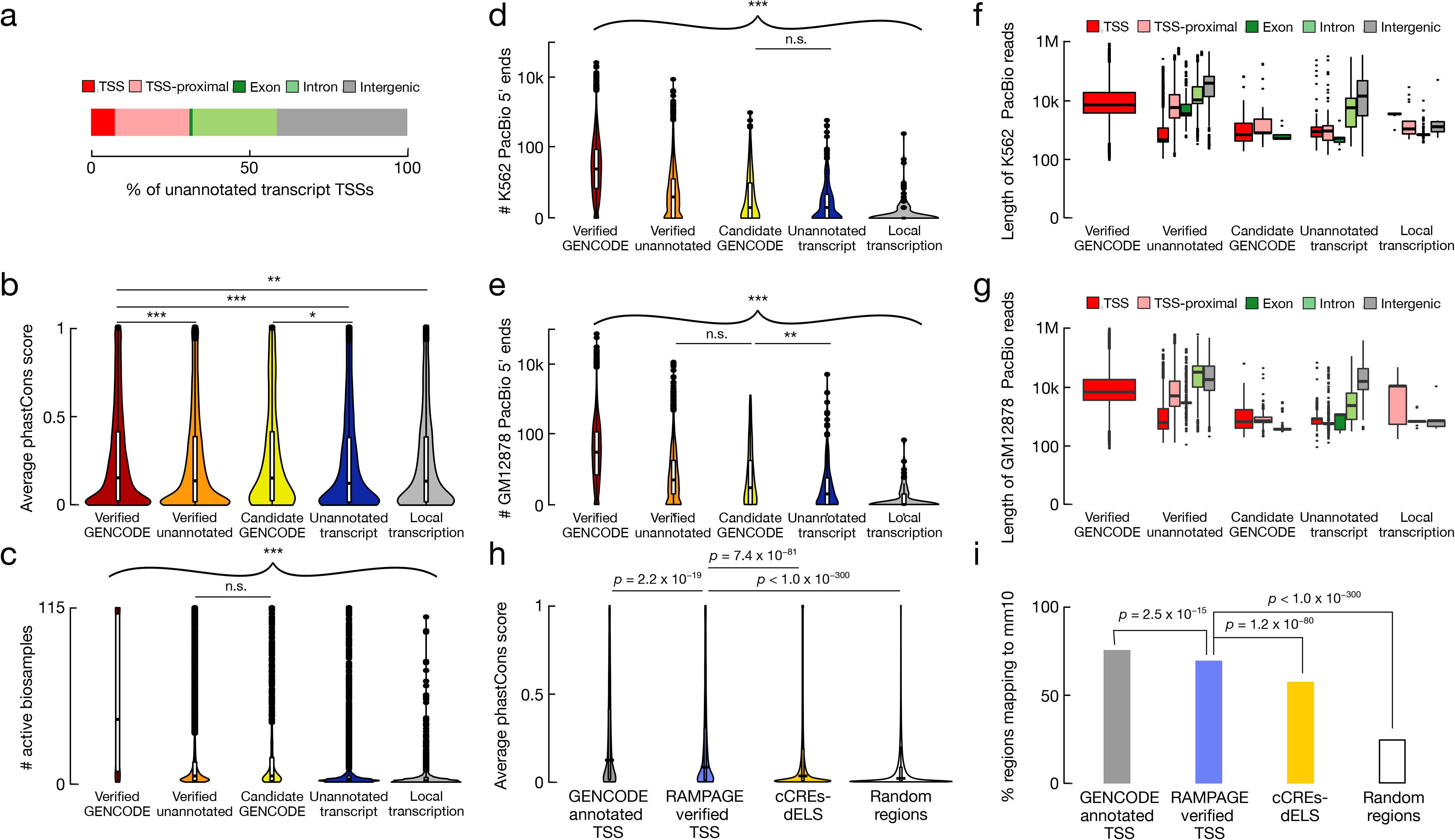
Features of RAMPAGE rPeaks stratified by TSS class. **a**, Bar graph displaying the genetic context of RAMPAGE rPeaks that were classified as TSSs of unannotated transcripts (TSS: red, TSS-proximal: pink, exon: dark green, intron: light green, intergenic: gray). **b,** Nested violin-boxplots displaying the average phastCons conservation score for RAMPAGE rPeaks stratified by TSS class (verified GENCODE: dark red, verified unannotated: orange, candidate GENCODE: yellow, unannotated transcript: dark blue, local transcription: gray). *P*-values corresponding to two-sided pairwise Wilcoxon tests with FDR correction are available in **Supplemental Table S4a**. Stars denote pairs with statistically significant differences (* *p* < 0.05, ** *p* < 0.01; *** *p* < 0.001) **c,** Nested violin-boxplots displaying the number of active biosamples (RPM > 2) of RAMPAGE rPeaks stratified by gene assignment category (as colored in **b**). *P*-values corresponding to two-sided pairwise Wilcoxon tests with FDR correction are available in **Supplemental Table S4c**. All pairs are statistically significant (*** *p* < 0.001) except where noted. **d,** Nested violin-boxplots displaying the number of overlapping K562 PacBio 5’ ends for K562 RAMPAGE rPeaks stratified by gene assignment category (as colored in **b**). *P*-values corresponding to two-sided pairwise Wilcoxon tests with FDR correction are available in **Supplemental Table S4c**. All pairs are statistically significant (*** *p* < 0.001) except where noted. **e,** Nested violin-boxplots displaying the number of overlapping GM12878 PacBio 5’ ends for GM12878 RAMPAGE rPeaks stratified by gene assignment category (as colored in **b**). *P*-values corresponding to two-sided pairwise Wilcoxon tests with FDR correction are available in **Supplemental Table S4d**. **f,** Boxplots displaying the length of K562 PacBio reads with overlapping 5’ ends for K562 RAMPAGE rPeaks stratified by gene assignment category and colored by genetic context (as defined in **a**). *P*-values corresponding to pairwise Fisher’s exact tests with FDR correction are available in **Supplemental Table S4e**. **g,** Boxplots displaying the length of GM12878 PacBio reads with overlapping 5’ ends for GM12878 RAMPAGE rPeaks stratified by gene assignment category and colored by genetic context (as defined in **a**). *P*-values corresponding to pairwise Fisher’s exact tests with FDR correction are available in **Supplemental Table S4f**. **h,** Nested violin-boxplots displaying the average phastCons conservation score for RAMPAGE-verified TSSs (purple), matched GENCODE-annotated TSSs (gray), cCREs-dELS (yellow), and random genomic regions (white). *P*-values correspond to two-sided pairwise Wilcoxon tests with FDR correction. **i,** Bar plots displaying the percentage of RAMPAGE-verified TSSs (purple), matched GENCODE-annotated TSSs (gray), cCREs-dELS (yellow), and random genomic regions (white) that liftOver to the mm10 genome. *P*-values correspond to pairwise Fisher’s exact tests with FDR correction.

